# Fast multi-resolution consensus clustering

**DOI:** 10.1101/2022.10.09.511493

**Authors:** Stijn van Dongen

**Affiliations:** Wellcome Sanger Institute, Hinxton, England (http://github.com/micans/)

**Keywords:** Multi-resolution clustering, network clustering, consensus clustering, Leiden, MCL, single-cell data

## Abstract

Clustering is a key tool for exploring results from large scale data collection efforts. It is especially useful to find clusterings (partitions of data into non-overlapping sets) at different levels of granularity, ranging from detailed to abstract. In order to be consistent such a set of clusterings should be nested, requiring that elements that group together in a given clustering group together in all coarser clusterings. Hierarchical clustering provides a tree for the data and naturally induces nested clusterings, but none of the methods have gained traction to provide such a multi-resolution view. Widely used methods such as Leiden and the Markov Cluster method (MCL) can produce clusterings at different scales controlled by a resolution parameter, but the results are generally not nested.

I introduce Restricted Contingency Linkage (RCL), a parameter-free consensus method that uniquely integrates and reconciles a set of flat clusterings with potentially widely varying levels of granularity into a single multi-resolution view. A common starting point in consensus methods is the association matrix *A*, where *A*_*ij*_ tallies the number of co-occurrences of *i* and *j* in the same cluster across all clusterings. RCL creates a matrix *R* where *R*_*ij*_ tallies a measure that differentiates *i*-*j* pairs with respect to the amount that they co-cluster in *pairs* of clusterings while taking into account the subset relationship between the clusters involved. The entries in *R* are fully abstracted from the input data, richly differentiated and uniformly normalised, making it suitable for single linkage clustering. From the resulting tree a limited set of flat clusterings is obtained by varying a resolution criterion along a logarithmic scale.

I validate the method using large-scale single-cell transcriptome data from human developing kidneys, along with visualisations and quality control measures of clustering ensembles. Marker gene expression is summarised in a heatmap across the multi-level RCL clusters, providing a coherent view of cell type and cell state hierarchies within the tissue. This reveals clusters corresponding to small populations of mast cells, plasma cells and interstitial cells, and shows segregation of cycling cells in small populations across different cell clades. Given an ensemble of input clusterings, RCL rapidly enables these analyses in a parameter-free manner.

An RCL reference implementation is provided for clustering ensembles that are associated with a network *G*, further restricting the RCL matrix to entries that correspond to edges in *G*. For a network *G* with *m* edges this implementation has complexity 𝒪(*m*(*p*^2^ + log(*m*))) where *p* is the number of input clusterings, taking less than a minute on a dataset with *N* = 27k elements, *m* = 1.5M edges and *p* = 24 clusterings. The implementation, including software for QC plots, marker gene heatmaps and other visualisations is available as part of the MCL software package (https://github.com/micans/mcl).

A suggestion for a quick guide to RCL is the pseudocode in Listing 1 and the multi-resolution cluster/marker-gene heatmap in Figures 11.1-11.2, supplemented with quality control plots for clustering granularity (Figures 8.1 and 8.1) and clustering discrepancy (Pages 5-6 and Figure 8.3).

## 1. Introduction

Clustering (a large class of methods) is a standard and often used approach in data analysis for separating data into groups, called clusters, often in large-scale high-dimensional data. A clustering (a data structure) is a partitioning of the data into disjoint clusters. This is sometimes called a *flat* clustering to explicitly distinguish from classification methods that yield a tree as outcome or from methods that allow overlapping clusters. Very broadly stated, the purpose of clustering is to find clusters such that the elements within a cluster are similar to each other and elements from different clusters are dissimilar. Several issues make this a challenging problem, two of which are especially of note. The first is that clustering is an underspecified problem. There is usually no model available for how groups may have formed in the data, what their (similarity) characteristics may be, or how many groups there may be. This makes it difficult to address the tension at the heart of clustering, namely that by grouping elements inevitably some between-cluster similarities of elements will be larger than some within-cluster similarities of elements. In the absence of a model for how groups have formed against the background of this trade-off, it is consequently difficult to establish a yardstick for the quality of a clustering that is principled and relevant for the data. The second issue is that cluster structure may be present and sought at different scales of resolution, reflecting that the notion of similarity is a matter of degree and perspective. Elements may be separated within one clustering but then group together within a second clustering where the element-wise similarity requirements are less stringent. In single-cell data this aspect is coming to the fore as datasets grow in size and complexity. Cell-type classifications exist at varying levels of specificity [1] and atlas data sets are commonly subclustered after clustering, often involving highly manual approaches [2, 3, 4, 5].

One long established way to address these issues is by employing one of the many hierarchical methods, examples of which are single link clustering [6], complete link clustering [7], upgma(unweighted pair group method with arithmetic mean) [8], and hierarchical clustering using Ward’s criterion [9]. The result of these methods is a tree; by cutting the branches of the tree a flat clustering can be obtained. For various reasons hierarchical methods are not prevalent in current large-scale data analyses such as employed for single-cell data. The methods are generally time consuming and do not scale well. Single linkage clustering is a relatively fast parameter-free method, but nearly always yields unbalanced clusterings due to a characteristic called *chaining*, where distant elements can cluster together due to a connecting chain of pairwise close elements.

A second way of addressing these issues is supplied by clustering methods that yield flat clusterings and have a resolution parameter that affects the scale of these. Examples of well established methods are Louvain [10], Leiden [11] and mcl [12]; the latter two are briefly described in Section 7. The challenge with these methods is that the clusters produced by each at different resolutions are generally not compatible: that is, the more fine-grained clusterings fail to subcluster the more coarse-grained clustering. This phenomenon is quantified and illustrated in Section 8. Hence, to find groupings at different levels manual approaches are used, such as subclustering parts of the data corresponding to individual clusters of interest [3, 4]. These supervised approaches can benefit from re-normalisation of data and careful guidance but are time-intensive. A fast parameter-free unsupervised multi-level clustering method may yield insights much sooner and can act as a map to the data for analysts from the start, from detailed to high-level structure.

I have had a long-standing interest in the discrepancies that exist between mcl clusterings derived with different inflation values (its resolution parameter). Initially this interest was focussed on the elements that cause these inconsistencies, those that switch allegiance and cluster with different sets of elements in different clusterings. Suppose an element *x* clusters in a cluster *A* in one clustering and a cluster *B* in another clustering. Eventually it dawned on me that the consistency of *x* is best seen as a continuous gradient that can be quantified by 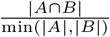 (the size of the intersection of *A* and *B* divided by the smaller of the size of *A* and the size of *B*). This measure yields one in those cases where *A* is a subset of *B* or vice versa, and it is close to zero if the size of the intersection |*A* ∩ *B*| is small relative to the sizes |*A*| and |*B*|.

It is a small step to then associate this measure with edges between element pairs that are in this intersection and tally the measure for all possible pairwise comparisons between clusterings. The resulting RCL matrix is a richly differentiated object that is nonetheless fully abstracted from the input data. It is also uniformly normalised, in the sense that each element is part of exactly one cluster in each clustering, and is part of exactly one intersection between clusters when comparing a pair of clusterings. This makes the RCL matrix suitable for single linkage clustering. Indeed, the issue of unbalanced clusters caused by so-called *chaining* does not arise as exemplified by the results in Section 8.

The result of applying single linkage to the RCL matrix is a hierarchical clustering in the form of a binary tree called the RCL tree. This tree has almost as many internal nodes as there are data elements.^1^ By picking a subset of internal nodes in the tree it is possible to obtain a flat clustering or a simplified hierarchical tree. There are many possible ways to do this, but all choices are compatible with the RCL tree and by extension with each other.

I propose a method to derive a flat clustering from the RCL tree by considering the deepest internal nodes that represent splits at a minimum level of resolution *r*. Varying this resolution threshold *r* on a logarithmic scale and taking all internal nodes corresponding to clusters in the clusterings thus obtained yields a simplified hierarchical view of the RCL tree. Associated with each cluster is the so-called *consistency level* at which it forms, giving a measure for its cohesiveness across the input ensemble. These procedures result in some proportion of nodes that are singletons or part of very small clusters. As a consequence of the RCL tree construction these nodes are the most ambiguously clustered data elements.

A short background of consensus clustering is provided in the next section. Following notations and definitions in Section 3, a small motivating example on a network of 16 nodes will be used to introduce both consistency and RCL in Section 4. Construction of the RCL matrix is formally described in Section 5. The application of RCL to the small example is then described in Section 6, with pseudocode for RCL provided on Page 11. The Leiden and mcl clustering methods are briefly described in Section 7. The final sections are mostly concerned with visualisation and validation of RCL results. Application of RCL is illustrated on a larger-scale dataset, introduced in Section 8, a filtered set of 27203 kidney cells [13]. Using this data the tree simplification algorithm is introduced in section Section 9. An alternative to umap plots is proposed in the form of a heatmap combining the RCL cluster hierarchy with a selection of marker genes (Section 11). Complexity and speed are summarised in Section 12, followed by discussion and acknowledgements.

## 2. Consensus clustering

*Consensus clustering* concerns the integration of multiple clusterings of the same dataset into a single result clustering [14], often called the consensus clustering or consensus partition. The same setting is described by many authors as the *cluster(ing) ensemble problem* [15]. In this and the majority of previous work cited, an input clustering is always a partition (sometimes called a flat or hard clustering) of the data into disjoint sets, as opposed to either a hierarchical clustering, an overlapping clustering, or a fuzzy clustering. I use the terms *clustering* and *partition* interchangeably, as well as the two terms *set* and *cluster*. To avoid excessive use of the word *clustering* the set of input clusterings is called the *input ensemble* and the resulting consensus clustering is called the *consensus outcome* in general, or the *consensus partition* or *consensus tree* depending on the nature of the outcome.

The clusterings in the input ensemble will have been computed on a set of entities and data describing these entities. Frequently this data is in the form of a numerical vector associated with each element, the vector representing measurements or scores on a number of features. In other cases the data may be in the form of a weighted network. Such a network can either be derived from feature data, such as gene expression in the case of single-cell data sets, or from direct one-to-one comparison of entities, as in the case of protein sequence similarity. The entities in the data are called *elements*. If these elements are represented in a network they are then also called *nodes*. Single linkage clustering is used to create a binary tree: the *leaf nodes* in this tree represent the data entities or elements, whereas an *internal node* in the tree represents a set or cluster of elements (namely all the leaf nodes below that node). Hence, entities, elements and nodes are loosely interchangeable in the setting of data, clusterings and networks; in the context of binary trees I will normally refer either to leaf nodes or internal nodes explicitly.

If the clusterings are derived from a network I call the input ensemble a *network ensemble*. A common choice or constraint is that the consensus outcome is computed without reference to the original data from which the input ensemble clusterings were derived, and this is largely the case here, except that in the specific case of a network ensemble the network can be used to speed up certain aspects of the computation.

Two different approaches can be distinguished in consensus clustering algorithms; these are the median/optimisation and the re-clustering/constructive approaches,^2^ with RCL belonging to the constructive class. In the median approach a clustering is sought that best represents the input ensemble, based on a criterion of proximity or distance between clusterings and a notion of optimal overall proximity between the consensus outcome and the input ensemble. The goal or expectation then is that the consensus partition should be an improvement on the input ensemble; the view is that of multi-learner improvement [15]. Examples of this approach use for example non-negative matrix factorization, genetic algorithms or kernels to seek or approximate the optimal solution.

In the constructive approach, the input ensemble is the starting point. The consensus outcome is newly built based on the input, often by applying another form of clustering to a data object derived from the input ensemble. For example, sc3 [17] performs *k*-means clustering several times to obtain the input ensemble. The weighted co-cluster matrix is clustered using hierarchical clustering with complete agglomeration and clusters are inferred at level *k* of the hierarchy. In both types of approach (median and constructive) previous work and reviews cited here are predominantly focused on consensus partitions as outcome. Consensus trees have been derived [17, 18] but were in this case seen as an intermediate step and accompanied with a strategy for deriving a partition from the tree.

## 3. Notation and definitions

Formulating RCL and generally considering input ensembles of partitions necessitates various comparisons of sets and partitions, quantification of these comparisons, as well a simple trait associated with binary trees. The required terminology and definitions are gathered in this section.

The input ensemble ℰ is a set of *p* partitions {𝒫_1_, …, 𝒫_*p*_}. The number of data elements is *d*. The elements are represented by the integers {1, …, *d*}, so that each 𝒫_*i*_ is a partition of {1, …, *d*}. This means that for each *i*, (i) 𝒫_*i*_ is a set of sets whose union is {1, …, *d*}, (ii) all of the pairwise intersections of sets in 𝒫_*i*_ are empty, and (iii) the empty set is not a member of 𝒫_*i*_. No further restrictions are imposed, in particular each partition may have its own number of constituent sets (clusters). A common starting point for consensus clustering is the association matrix defined below.

Definition 3.1. *The association matrix A for the input ensemble* ℰ *has d rows and d columns. The value A*_*xy*_ *in row x and column y is defined as follows*.

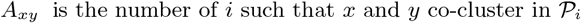

From the definition it follows that *A* is symmetric, that is, *A*_*xy*_ = *A*_*yx*_ for all *x* and *y*. The single-linkage hierarchy derived from an association matrix will be used in a small motivating example.

RCL uses a criterion called *set consistency* for how close either of the sets is to being a subset of the other.

Definition 3.2. *The set consistency* scy(*A, B*) *of two sets A and B is zero if one or both sets are empty. If both sets are non-empty it is defined by*

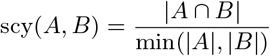

If both sets are non-empty and one is a subset of the other then scy(*A, B*) equals 1. If the two sets have empty intersection it equals 0. For example, the consistency of the two sets {1 2 3 4 5} and {3 4 5 6} is 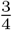. Consistency is known in the literature as the *overlap coefficient* or the *Szymkiewicz-Simpson coefficient*. I use *set consistency* here as that term has a meaning that fully aligns with the purpose of measuring cluster agreement.

Given two clusterings 𝒫 and 𝒬 of {1, …, *d*}, of interest is the clustering that is called their meet or intersection, or more precisely the greatest common subclustering.

Definition 3.3.

*The greatest common subclustering of* 𝒫 *and* 𝒬 *is denoted* gcs(𝒫, 𝒬) *and obtained by taking all non-empty intersections A* ∩ *B where A, B range over all sets in* 𝒫 *and* 𝒬, *respectively*.

The greatest common subclustering of 𝒫 and 𝒬 is useful to describe between-clustering discrepancies. This applies both to (clusterings obtained from) a single input ensemble and multiple input ensembles. In practice most input ensembles will have clusterings that do not nest. That is, when comparing pairs of clusterings from among the ensemble it will usually not be the case that one is a perfect subclustering of the other. It is useful to quantify *how close* a clustering is to being a subclustering of another. The dpy function below describes this as a discrepancy fraction, that is, the fraction of elements that needs to be split off from one clustering to obtain a subclustering of the other clustering. A pair of these scores can be used to gauge the relationship between two clusterings.

Definition 3.4. *For partitions* 𝒫 *and* 𝒬 *of* {1, …, *d*} *the discrepancy* dpy _𝒬_ (𝒫) *of* 𝒫 *with respect to* 𝒬 *is defined as*

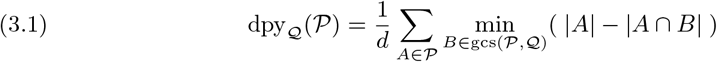

This number can be interpreted as the difference between 𝒫 and the greatest common sub-clustering of 𝒫 and 𝒬 as a fraction of the number of elements.^3^ For each cluster *A* in 𝒫 the quantity min_*B*∈gcs(𝒫, 𝒬)_(|*A*| − |*A* ∩ *B*|) is the smallest number of elements from *A* such that *A* with those elements removed yields a set in gcs(𝒫, 𝒬). This is equivalent to picking the best match of *A* in gcs(𝒫, 𝒬), the cost being the number of elements in *A* not present in that best match. If the sum of these costs equals 0 then 𝒫 is identical to gcs(𝒫, 𝒬) (each cluster in 𝒫 is also a cluster in gcs(𝒫, 𝒬)) and thus 𝒫 is a subclustering of 𝒬.

Discrepancy between all pairs of clustering from an input ensemble can be plotted in a heatmap to visualise this aspect of the internal structure of the ensemble, as seen below.

**Partition discrepancy example** Four partitions 𝒫, 𝒬, ℛ, 𝒯 and their six greatest common subclusterings are given below.

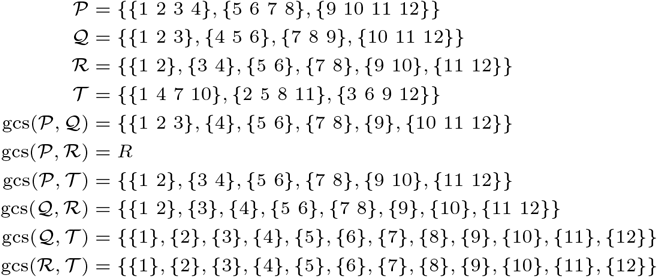

The discrepancies derived from these are tabulated below. A second presentation as heatmap is given as well; this form will be used to visualise Leiden and mcl discrepancies.

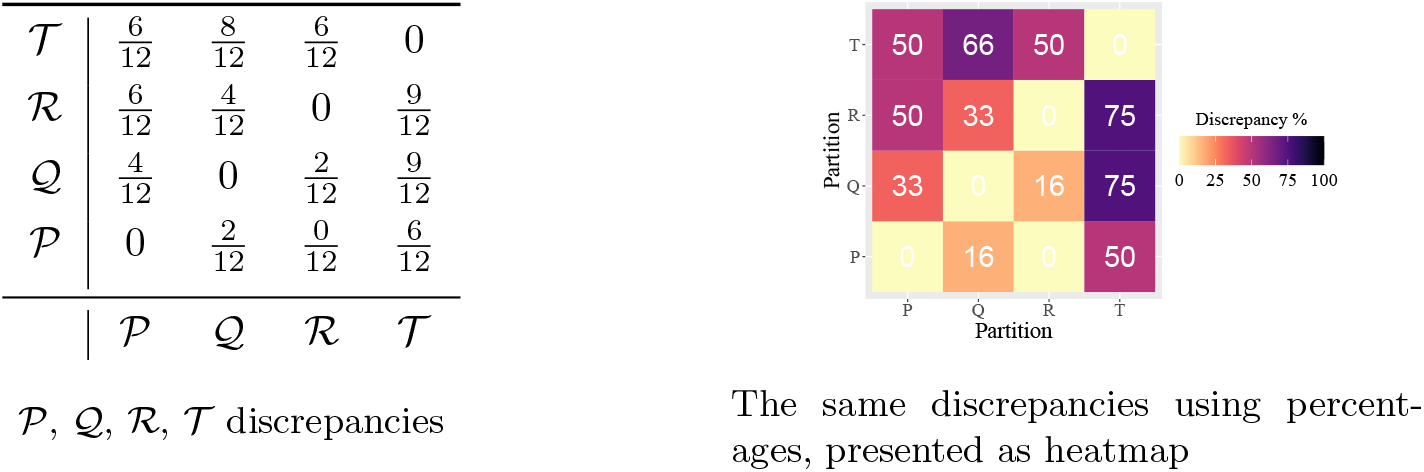

To define RCL no notation is needed or provided to index a partition to list its constituent sets or the elements within those sets. However, it is useful to establish the reverse, to find for a pair of elements *x, y* and a partition 𝒫 whether *x* and *y* reside in the same cluster in 𝒫 (that is, they co-cluster). To this end I introduce the set-finding function **S**.

Definition 3.5. *For a partition* 𝒫 *and two elements x, y, the set-finding function*

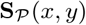

*yields the empty set if x and y do not co-cluster in* 𝒫 *and otherwise the cluster of which they are both part*.

For the RCL tree simplification algorithm I will use the notion of *largest sub-split* or lss. Any node *n* in a binary tree is associated with a subtree of which it is the root. The size of that subtree rooted at *n* is written |*n*|. It is the count of all the leaf nodes below *n*, or 1 if *n* is a leaf node itself.

Definition 3.6. *The split of an internal node n in a binary tree is the size of the smallest of the two subtrees associated with the two children of n. The largest sub-split* lss(*n*) *associated with n is the largest split among n or any of its descendants*.

When building a binary tree lss can be easily computed for each node as the tree is built. When merging nodes *x* and *y* in a parent node *z*, the largest sub-split for *z* can be computed as

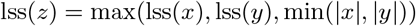

## 4. Small example

The starting point of RCL is the observation that within an input ensemble there is in general variability in how consistently elements cluster and co-cluster. Consider Figure 4.1. It depicts a grid-like network with 16 nodes and 24 edges, with three partitions 𝒫_1_, 𝒫_2_, 𝒫_3_. Cluster membership is indicated by node colour.^4^ It was already stated that RCL is expected to be most useful for input ensembles of varying granularity. This example does not feature such variability, moreover all 𝒫_*i*_ have the same number of clusters. However, the principle informing RCL tree construction is equally well explained in the absence of such variability. The multi-resolution aspects of RCL will subsequently be introduced using input ensembles defined on a large-scale single-cell kidney data set [13].

**Fig. 4.1:**
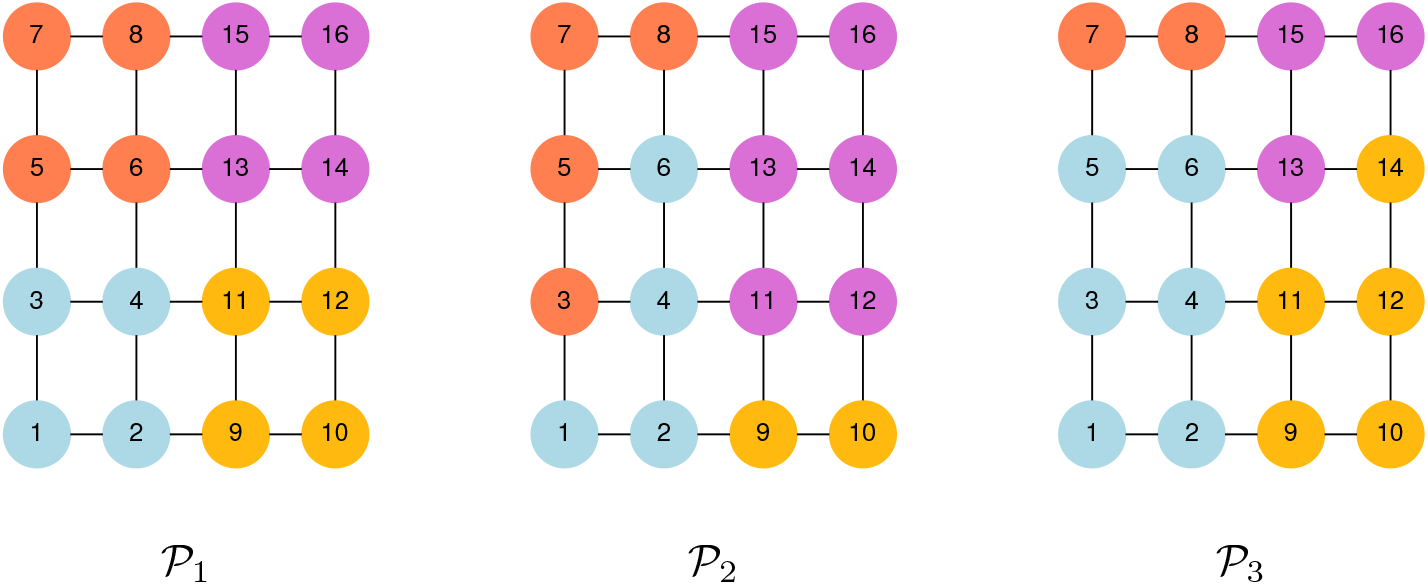
A network with three different clusterings, clusters indicated by the node colours.

The three partitions were artificially imposed on the network. Nonetheless, the example is not unrealistic, as similar shifts of allegiance do happen in practice. One can imagine that the differences are a result of different algorithms, perhaps with randomised aspects, or influenced by edge weight differences not depicted here. In any case, this is the example that will illustrate the principle of node clustering consistency. The three partitions are again presented below as lists of sets.

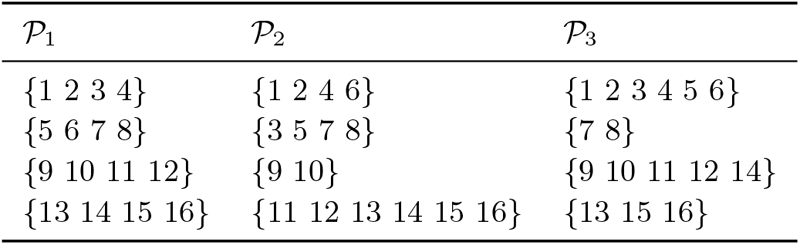

| $\mathcal{P}_1$ | $\mathcal{P}_2$ | $\mathcal{P}_3$ |
| --- | --- | --- |
| $\{1\ 2\ 3\ 4\}$ | $\{1\ 2\ 4\ 6\}$ | $\{1\ 2\ 3\ 4\ 5\ 6\}$ |
| $\{5\ 6\ 7\ 8\}$ | $\{3\ 5\ 7\ 8\}$ | $\{7\ 8\}$ |
| $\{9\ 10\ 11\ 12\}$ | $\{9\ 10\}$ | $\{9\ 10\ 11\ 12\ 14\}$ |
| $\{13\ 14\ 15\ 16\}$ | $\{11\ 12\ 13\ 14\ 15\ 16\}$ | $\{13\ 15\ 16\}$ |

Following [18], simply treating the association matrix *A* (not shown here) as a similarity matrix, the single linkage tree associated with *A* is shown in Figure 4.2.

**Fig. 4.2:**
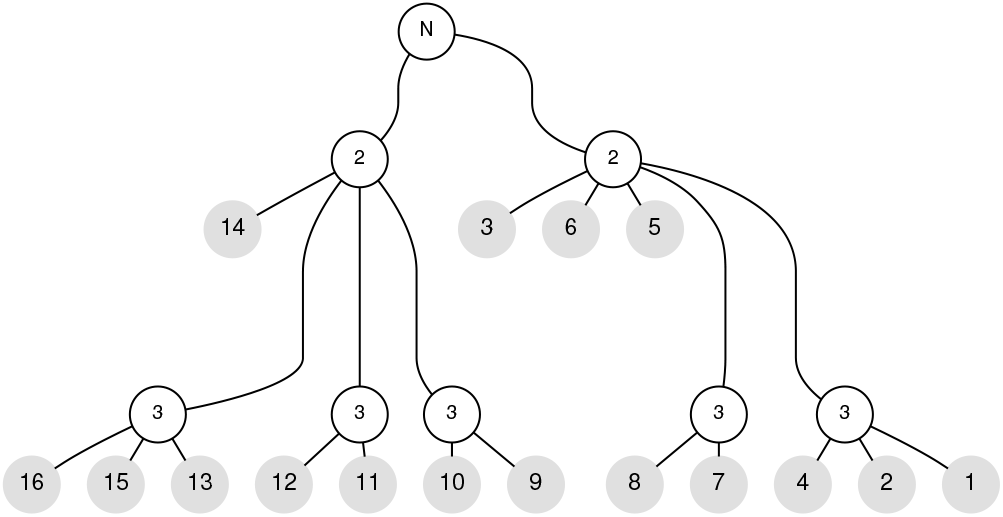
The single linkage tree derived from the association matrix *A* for the clusterings in Figure 4.1. Grey nodes are leaf nodes. Internal nodes are labelled with the co-clustering value from *A* at which all the nodes below are transitively linked. The root labelled *N* indicates the final join is based on network connectivity.

In this article network edge weights and matrix entries always represent similarities rather than distances. Single linkage clustering thus considers node pairs in order of highest similarity rather than smallest distance.

The tree in Figure 4.2 reflects commonality of clustering and the transitive property of single linkage clustering. That is, if node *x* co-clusters twice with *y* and *y* co-clusters twice with *z*, then all three merge at the same time. The clusters that are determined by the internal nodes of the tree at level 3 are two triplets, four pairs and four singletons, and at level 2 are two sets of eight nodes. It is easy to verify for example from Figure 4.1 or the table above that {1 2 4} and {13 15 16} co-cluster in all three partitions. Additionally it can be verified that the late-joining nodes 3, 5, 6 and 14 co-cluster twice with some nodes but never more often than that.

Consider the following claim: The co-clustering twice of nodes 3 and 5 is of a more tenuous nature than the twice co-clustering of 9 and 11. Observe for example that the latter two nodes are part of the larger group of nodes {9 10 11 12} that co-clusters in its entirety in both 𝒫_1_ and 𝒫_3_. Loosely speaking the nodes 3 and 5 seem in limbo between the areas in the top left and the bottom left of the network. This can be made more precise by considering intersections between clusters.

The pair of elements 3, 5 forms the entire intersection of the clusters *A* = {3 5 7 8} in P_1_ and *B* = {1 2 3 4 5 6} in P_3_, respectively, noting that 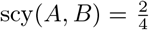. This pair co-clusters with quite different sets of nodes when considering these two partitions. The pair (9, 11) is part of the larger intersection of the clusters *C* = {9 10 11 12} and *D* = {9 10 11 12 14}, where 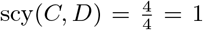. The intersections tell us something about nodes and clustering consistency. Node 3 is the sole inhabitant of the intersection of the clusters *A* = {1 2 3 4} and *B* = {3 5 7 8} in P_1_ and P_2_. This makes 3 a volatile presence and detracts from the claim that 3 is a good co-clusterer in either of these sets. This is quantified by scy 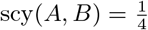.

The above is an illustration that set consistency is potentially useful in differentiating between different co-clustering scenarios. If *A* is a subset of *B* or *B* is a subset of *A* then they are in full agreement and their classification of elements fits into a multi-resolution view. The approach taken in RCL is to quantify consistency in a refined version of the input ensemble association matrix.

## 5. RCL description

The restricted contingency matrix *R* is defined in terms of sums of consistency scores (defined in Definition 3.2) between sets. The explanation following Formula 5.1 and the pseudocode in Listing 1 provide perhaps the easiest ways to understand the following definition.

Definition 5.1. *For an input ensemble* ℰ *of clusterings* {𝒫_1_, …, 𝒫_*p*_}, *the associated restricted contingency matrix R is defined by setting the xy entry*

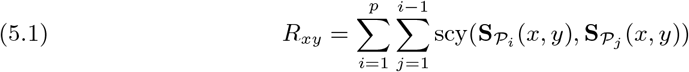

*Equivalently, R*_*xy*_ *is the sum of the consistency scores for every pair of clusters taken from pairs of distinct partitions in the input ensemble where x and y co-cluster in both partitions*.

*For a network G and an input ensemble of clusterings, the associated restricted contingency matrix R*_*G*_ *is defined by setting its xy entry to R*_*xy*_ *if* (*x, y*) *form an edge in G, and to zero otherwise*.

One can think of the concurrent co-clustering mandated by Definition 5.1 as *weighted co-co-occurrence*. For two elements *x* and *y* the sum in Equation 5.1 is over pairs of partitions 𝒫_*i*_ and 𝒫_*j*_ where *x* and *y* co-cluster in both,^5^ the summand being the consistency of the respec-tive 𝒫_*i*_ and 𝒫_*j*_ clusters in which they are both contained. The maximum value that *R*_xy_ can assume is the number of comparisons made, that is, 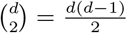. This will be the case if *x* and *y* always co-cluster in all partitions and every intersection where *x* and *y* occur has consistency 1.

Definition 5.2. *The* RCL *tree for an ensemble* ℰ *is defined as the tree obtained by applying single linkage clustering to the restricted contingency matrix R*.

For network ensembles, a useful optimisation method is to use the connectivity structure of *G* to limit the number of entries of the restricted contingency matrix. Both *R* and *R*_*G*_ should yield the same tree when applying single linkage clustering in all reasonable cases.^6^ An implementation of RCL should not use Definition 5.1 directly. It is more efficient to iterate over all unordered pairs of partitions 𝒫_*i*_, 𝒫_*j*_, then iterate over all intersections in gcs(𝒫_*i*_, 𝒫_*j*_), as illustrated in the pseudocode in Listing 1. For network ensembles one can update edges in *R*_*G*_ that have both source and sink in the intersection, otherwise one can update *R*_*xy*_ for all pairs of *x, y* in the intersection.

The term consistency is used with two meanings. The first is the consistency between two sets as in Definition 3.2. The summed and scaled consistency scores in Definition 5.1 and Definition 5.2 are also referred to as consistency; it should be clear from the context which meaning is intended. The reference implementation rescales the consistency values in Definition 5.1 to lie between zero and one thousand, that is, the value between zero and one is expressed as an integer count of thousandths. This scaling is used in subsequent figures where consistency is one of the dimensions or traits.

The Clustree method [20], a tool for visualising and evaluating clusterings at multiple resolutions, uses the contingency structure between clusterings to establish a lattice-type graph between clusterings in an input ensemble. The clusterings are ordered from coarse to finegrained; for our purposes, assume that the partitions 𝒫_*i*_ are ordered in this manner. Clustree establishes a link between a cluster *A* in 𝒫_*i*_ and a cluster *B* in 𝒫_*i*+1_ if their intersection is non-empty, with a weight called *in-proportion* set to 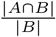. This results in a lattice representation on the clusters of the clusterings in the input ensemble, providing *a compact, information-dense visualization that can display summarized information across a range of clusters* [20]. Given the Clustree ensemble ordering, one may generally expect the second set *B* to be smaller in size than *A*. Hence, in-proportion will in general be identical to the set consistency used by RCL. There is a conceptual link between Clustree and RCL in that they both utilise the contingency structure of the input ensemble and use a similar weight measure. Clustree utilises this weight to visualise and summarise clustering to clustering and cluster to cluster overlap, whereas RCL pulls the same information down to the level of node-node co-co-occurrence, tallies it, and thus obtains a refinement of the association matrix suitable for single-linkage clustering.

## 6. Small example with RCL applied

The table below lists, for the previous example, all of the non-empty pairwise intersections between clusters from different partitions, the size of the two clusters that give rise to each of the intersections, and the consistency of the intersection. Fractions have not been simplified to emphasize the link to the listed set sizes.

| $\text{gcs}(\mathcal{P}_1, \mathcal{P}_2)$ | scy | $(\vert S_1\vert , \vert S_2\vert )$ | $\text{gcs}(\mathcal{P}_1, \mathcal{P}_3)$ | scy | $(\vert S_1\vert , \vert S_3\vert )$ | $\text{gcs}(\mathcal{P}_2, \mathcal{P}_3)$ | scy | $(\vert S_2\vert , \vert S_3\vert )$ |
| --- | --- | --- | --- | --- | --- | --- | --- | --- |
| {1 2 4} | 3/4 | (4,4) | {1 2 3 4} | 4/4 | (4,6) | {1 2 4 6} | 4/4 | (4,6) |
| {3} | 1/4 | (4,4) | {5 6} | 2/4 | (4,6) | {3 5} | 2/4 | (4,6) |
| {5 7 8} | 3/4 | (4,4) | {7 8} | 2/2 | (4,2) | {7 8} | 2/2 | (4,2) |
| {6} | 1/4 | (4,4) | {9 10 11 12} | 4/4 | (4,5) | {9 10} | 2/2 | (2,5) |
| {9 10} | 2/2 | (4,2) | {14} | 1/4 | (4,5) | {11 12 14} | 3/5 | (6,5) |
| {11 12} | 2/4 | (4,6) | {13 15 16} | 3/3 | (4,3) | {13 15 16} | 3/3 | (6,3) |
| {13 14 15 16} | 4/4 | (4,6) |  |  |  |  |  |  |

<sup>6</sup>That is, counter-examples may exist but are likely to be tortuous and artificial, as it would require two nodes not connected in $G$ to co-cluster directly before being joined transitively by co-clustering of the nodes on a path between them.

### Listing 1

Restricted Contingency Linkage pseudocode. In the presence of a network *G* the shaded code can be used to optimise the implementation. The last step, simplification, is described in Section 9.

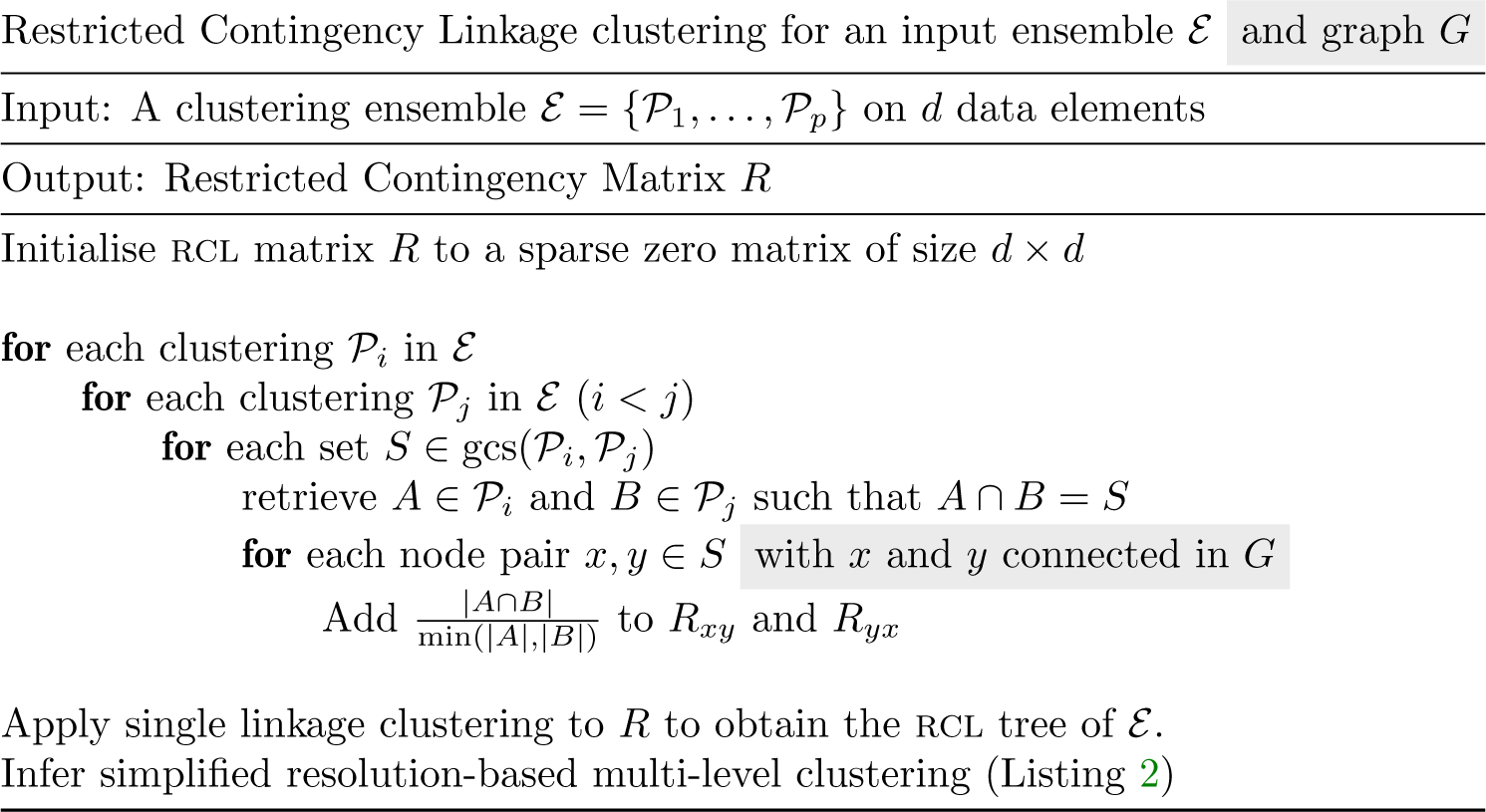

Using this table it can be seen for example that 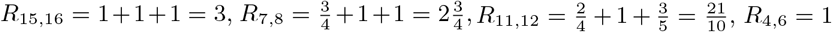. Computation of all values leads to the tree in Figure 6.1. In the figure the values in the internal nodes are the summed values as in Definition 5.1, subsequently scaled between 0 and 1000, following the convention in the implementation used. In this example, the maximum RCL value attainable is three, as there are three possible comparisons between the three partitions. Scaling to 1000 requires a scaling factor 1000*/*3, so that in the figure nodes 11 and 12 joint at a value of 1000·*R*_11,12_*/*3 = (21·1000)*/*(10·3) = 700. These summed and scaled values are called the (RCL) *consistency values*.

**Fig. 6.1:**
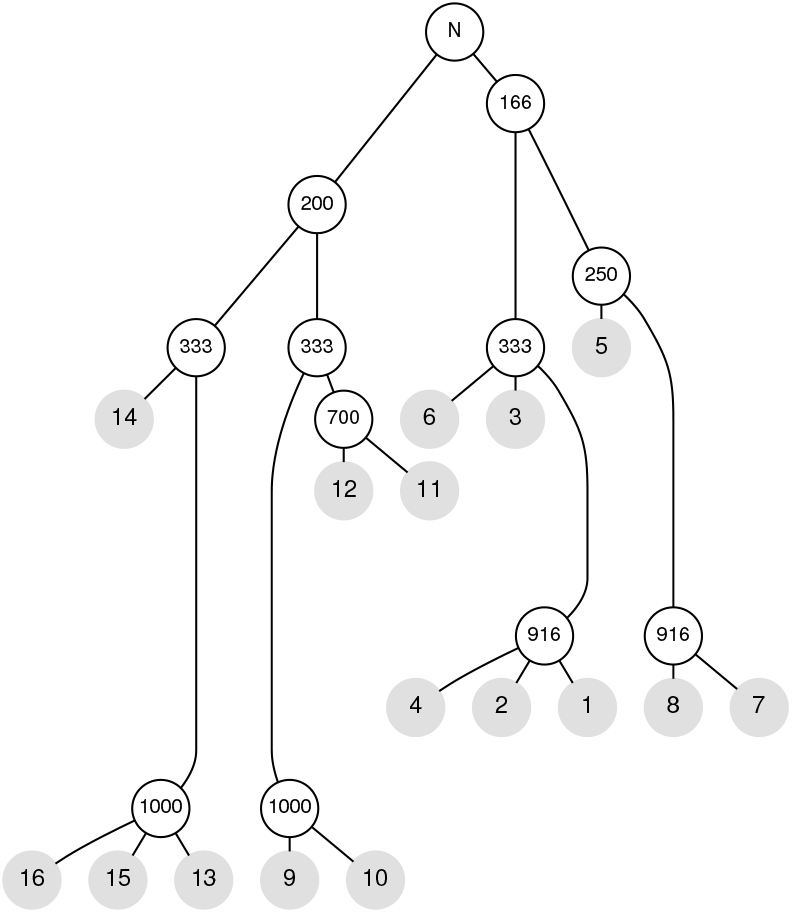
The RCL tree constructed for the clusterings in Figure 4.1. Internal nodes are labelled with the co-clustering value from the RCL matrix at which all the nodes below are transitively linked. Values are rescaled by a factor 1000*/*3 so that the maximum value is 1000. The root labelled *N* indicates the final join was based on network connectivity.

The tree in Figure 6.1 has, unsurprisingly, a more intricate structure when compared to the tree in Figure 4.2. The tree has internal nodes that correspond roughly to the four quadrants of the network, with {9 10 11 12}, {13 14 15 16}, and {1 2 3 4 6} all grouping together at tree consistency level 333. The nodes {5 7 8} group later at level 250, reflecting the unconvincing and volatile character of node 5 described above. The groups {13 15 16} and {9 10} both form at the highest possible consistency level, indicated that they only ever appear in fully consistent cluster pairs, where one is always a subset of the other. The group {1 2 4}, although always co-clustering together, appears in one intersection of lesser consistency (between 𝒫_1_ and 𝒫_2_) and hence forms at a slightly lower level.

In this tiny example the RCL matrix, a refined version of the cluster ensemble association matrix, looks to have some merit. In order to consider large-scale data, a way is now needed to simplify and interpret the single linkage hierarchy obtained from the RCL matrix. This is described in Section 9. The Leiden and mcl methods are described first in Section 7 and the kidney data set is introduced in Section 8.

## 7. The Leiden and mcl clustering methods

Leiden [11] and mcl [12] are both clustering algorithms that produce flat clusterings and have a single parameter to control cluster granularity. Leiden is based on optimisation of *modularity* [21, 22] and falls within a class of modularity optimisation approaches including FastModularity [23] and Louvain [10]. Leiden has a resolution parameter *γ* that controls the granularity of the output clustering. It does so indirectly, without specifying the number of clusters. Lower values of *γ* lead to larger and fewer clusters. The Leiden algorithm is widely used and part of the standard Seurat [24] and Scanpy [25] workflows for single-cell data analysis.

MCL is based on computing random walk probabilities through a network, alternating the computation of two-step probabilities with a step called *inflation*. Inflation pushes probabilities apart, causing larger probabilities to dominate lower probabilities. This step is parameterised by the inflation parameter, with lower values again leading to larger and fewer clusters. The mcl algorithm is widely used in protein sequence clustering [26, 27, 28] and other application areas such as protein interaction networks [29], gene expression networks [30] and functional interaction networks based on association data [31].

Here I use both Leiden and mcl on the kidney data set described in the next section. For each method an input ensemble of 12 clusterings on this data set is generated. The ensembles are quantitatively described in terms of cluster granularity and both their internal and mutual partition discrepancies.

## 8. Single-cell transcriptomics of developing human kidneys

A large-scale RCL analysis was performed using a kidney data set [13, 32, 33], briefly described below. This section further consists of a quantitative description of two clustering ensembles for this data, for Leiden and mcl respectively. The RCL tree simplification algorithm is described in the next section.

For the kidney data a filtered set with 27203 cells from developing human kidneys was used [13]. This was processed with a standard Seurat workflow. Cell/gene data was embedded in a 50-dimensional space, from which shared nearest neighbour networks were derived based on *k*-NN networks with *k* = 20 for Leiden and *k* = 30 for mcl. For mcl a higher value was used to nudge it towards coarser clusterings. Allowing more edges results in slightly larger clusters, although the effect is not pronounced.

Leiden [11] clustering was performed using twelve resolution parameters 0.6, 1, 1.4, 2, 4, 6, 8, 10, 12, 14, 16 and 20. mcl clustering was performed using twelve inflation parameters 1.3, 1.35, 1.4, 1.45, 1.5, 1.55, 1.6, 1.65, 1.7, 1.8, 1.9 and 2. For mcl the granularity of results has a power-type dependency on the inflation parameter, hence its increased sampling rate at the lower end. For Leiden the set of parameters was similarly chosen so that the granularity characteristics of the clusterings were somewhat evenly spread throughout the Leiden input ensemble. It can be noted that the first three Leiden resolution parameters (0.6, 1, and 1.4) fall roughly in the range recommended for Seurat and Scanpy, leading to large clusters. At the other end, 20 is a little bit below the value 25 used by Celltypist [1] to oveRCLuster data in order to allow for fine-grained cell-type assignment.

Both sets of parameters were selected after confirming the spread of cluster sizes using the granularity signatures as displayed in Figures 8.1 and 8.2 and the clustering discrepancies in Figure 8.3. The reference RCL implementation (Section 13) provides the program RCL-qc to create these plots as a quality control step. The granularity signatures show for each resolution and inflation value the proportion of elements in a cluster of size at most *x* as *x* is varied. The shape of the curves are similar across both methods as well as their varying parameters. The curves for Leiden are shifted a bit further to the right, indicating that the Leiden ensemble as a whole has slightly coarser clusterings when compared to the mcl ensemble. It is beyond the scope of the current article to investigate optimisation of selection of the input clusterings, but it seems reasonable to aim for separation and even distribution of the granularity signatures such as seen in Figures 8.1 and 8.2. This is both to promote a multi-level view and to avoid skewed contributions due to very similar clusterings, as might be the case if two or more such signatures were to clump together.

**Fig. 8.1:**
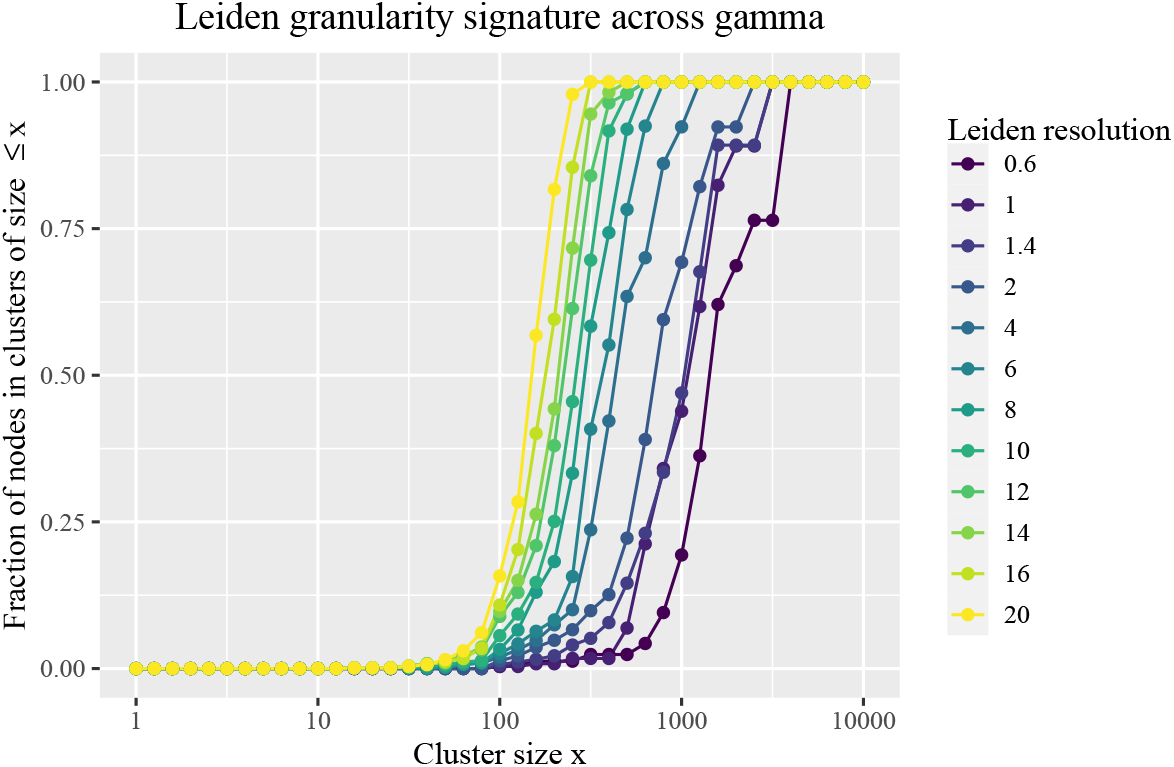
Each graph corresponds to a clustering of the kidney data (Section 8) for the specified Leiden resolution value (also called gamma). The graphs show the fraction of elements that are part of a cluster of size at most *x* for varying *x*.

**Fig. 8.2:**
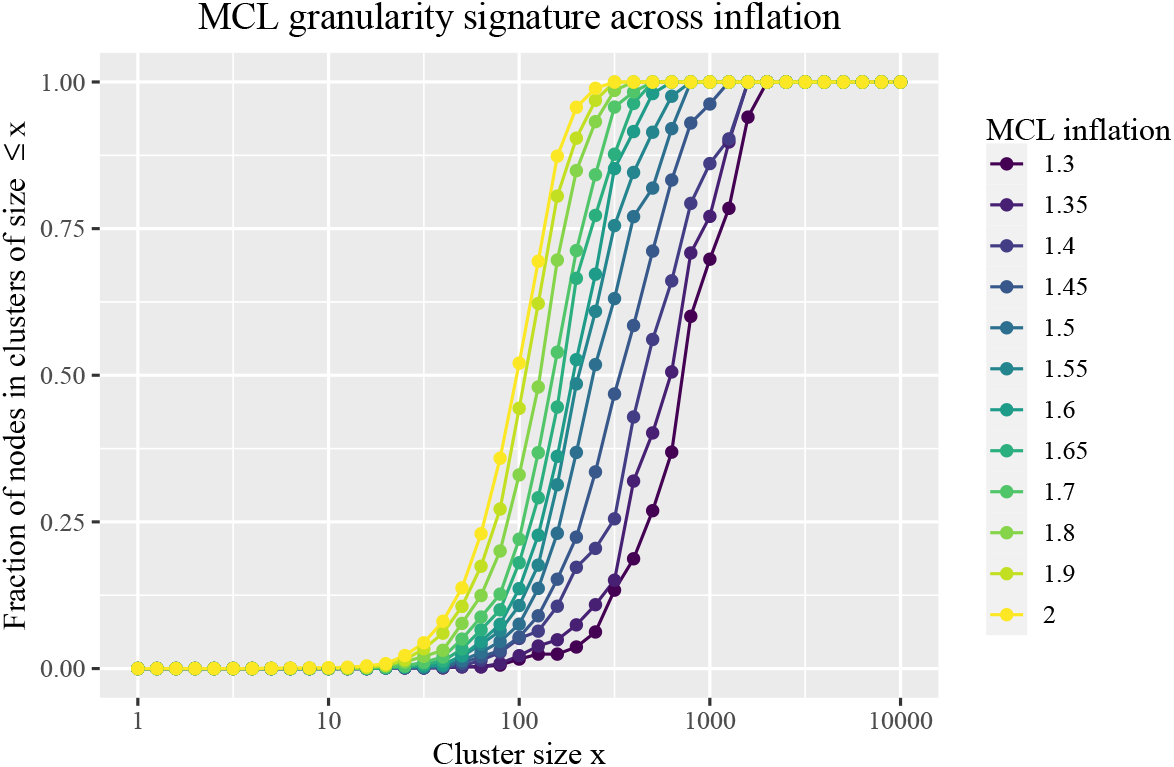
Each graph corresponds to a clustering of the kidney data (Section 8) for the specified mcl inflation value. The graphs show the fraction of elements that are part of a cluster of size at most *x* for varying *x*.

**Fig. 8.3:**
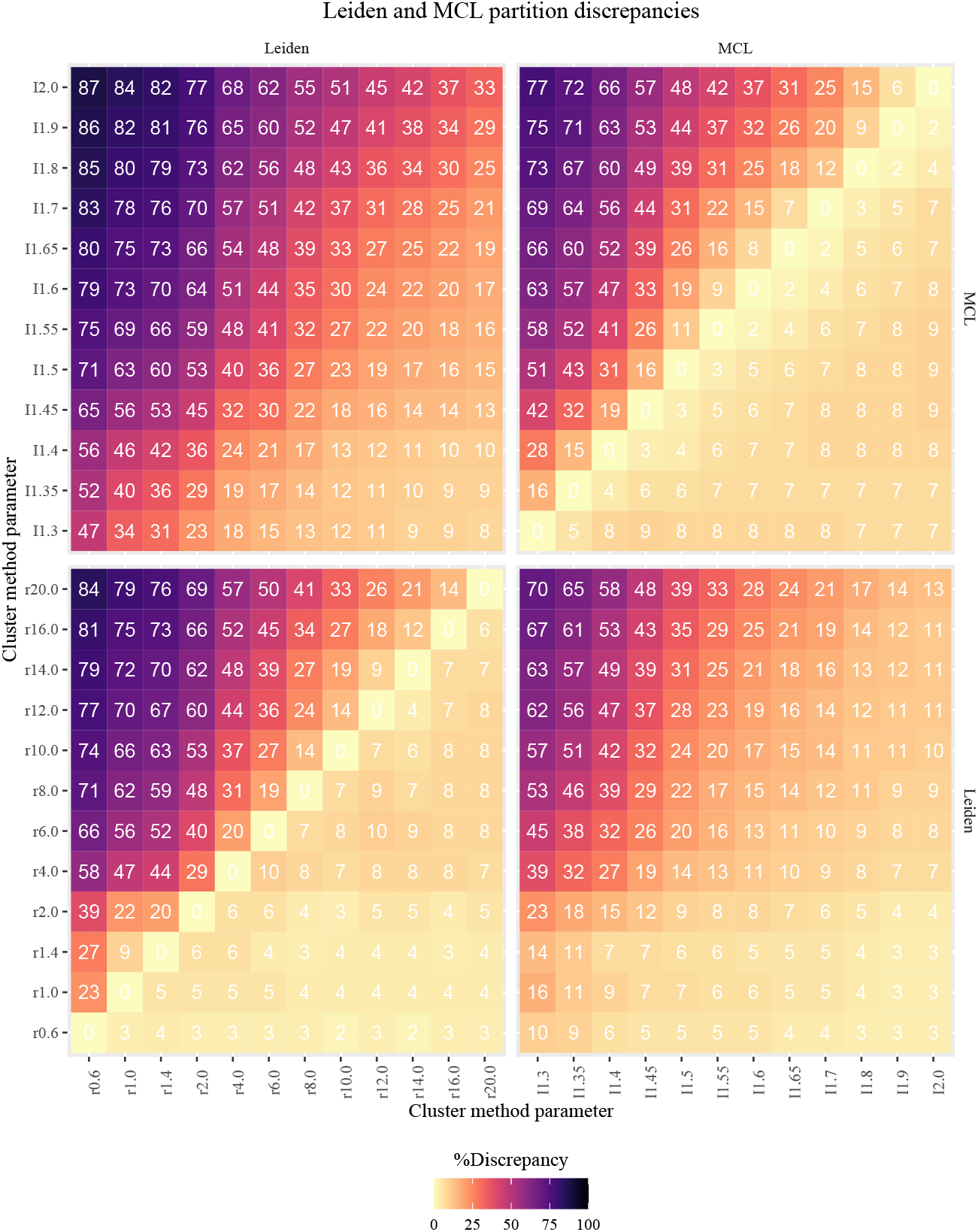
Partition discrepancy (expressed as percentage) for Leiden and mcl clusterings. The *i, j* value denotes the percentage of elements needed to split off in order to transform 𝒫_*i*_ into gcs(𝒫_*i*_, 𝒫_*j*_). Low *i, j* value compared to *j, i* implies that 𝒫_*i*_ is close to being a subclustering of 𝒫_*j*_.

The granularity plot shows the distributions of cluster sizes but provides no information about the contingency (intersection) structure of clusterings in the ensemble. Complementary to this, the heatmap of clustering discrepancies, introduced in Section 3, provides no information about cluster sizes but summarises the contingency structure between clusterings, here shown in Figure 8.3 for the Leiden and mcl kidney data clustering ensembles. These discrepancies unambiguously establish whether clusterings are very similar, the extent to which subclustering is present, or if two or more groups in the clustering ensemble represent conflicting views, as was the case with clustering 𝒯 versus clusterings 𝒫, 𝒬, and ℛ on Page 6. Additionally the heat map is informative for how evenly placed apart the clusterings are in terms of discrepancies.

The plot shows cluster discrepancies for pairs of clusterings chosen from all pairings between the Leiden and mcl ensembles, logically separated into Leiden-Leiden, mcl-mcl, Leiden-mcl and mcl-Leiden quadrants. The most important part in each quadrant is the lower right, where its smallest values are found. For the Leiden-Leiden and mcl-mcl quadrants the lower right always contains the clustering discrepancy from a finer-grained clustering with respect to a coarser-grained clustering. The values in these parts indicate the percentage of elements needed to transform the former into a subclustering of the latter, *or equivalently, how close the former is to being a subclustering of the latter*.

The median of the values in the lower right Leiden-Leiden quadrant is 5, for mcl it is 7. This indicates that for Leiden and mcl respectively, on average 5 and 7 percent of elements need rearranging to transform 𝒫 into a subclustering of 𝒬. In my experience, such values are typical when analysing large-scale single-cell data. Their spread and range can be considered a baseline when analysing other data or using other methods.

The median values for the upper left parts of the same quadrants are 46 for Leiden, and 38 for mcl. This signifies that 𝒫 and 𝒬 are a bit further apart on average for Leiden as compared to mcl, and reflects the tendency towards coarser Leiden clusterings as already observed in the granularity signatures. Additionally the heatmap shows a smooth distribution of discrepancies for both the Leiden and mcl ensembles as their respective parameters are varied. There are no sudden jumps in the heat map values as one moves from a clustering at one resolution or inflation value to the next. Finally, the heat map also shows the level of agreement between (clusterings from) the Leiden and mcl input ensembles. The smallest values (those on the lower right of the quadrants for respectively the Leiden-mcl and the mcl-Leiden comparisons) are a little bit larger than the corresponding values in the Leiden-Leiden and mcl-mcl quadrants, indicating a lesser amount of subclustering. This is to be expected given that the methods operate on quite different principles; there is certainly no evidence of outright divergence or disagreement.

## 9. Characteristics of the RCL tree and a tree simplification algorithm

The RCL single linkage hierarchy for the kidney data is a binary tree with 27203 leaf nodes and 27202 internal nodes for both the Leiden and mcl input ensembles, integrating all the data points into a single tree. In this case this was due to the implementation making use of the underlying network connectivity and the network consisting of a single connected component. In other instances the RCL outcome may be a collection of distinct trees. Equally an implementation may have an option to artificially add a root partition with a single cluster containing all data elements to always ensure a single tree as outcome. It is of interest to consider the distribution of consistency values associated with the internal nodes of an RCL tree for an input ensemble. This is shown for both the Leiden and mcl ensembles in Figure 9.1. Both distributions show the large bulk of values to be between 750 and 1000, with the largest part of the remaining tail between 500 and 750. These plots show the differentiation and spread in edge weights in the RCL matrix and tree, contrasting with the maximum amount of twelve different nonzero values that could potentially occur in the association matrix for an input ensemble with twelve clusterings.

**Fig. 9.1:**
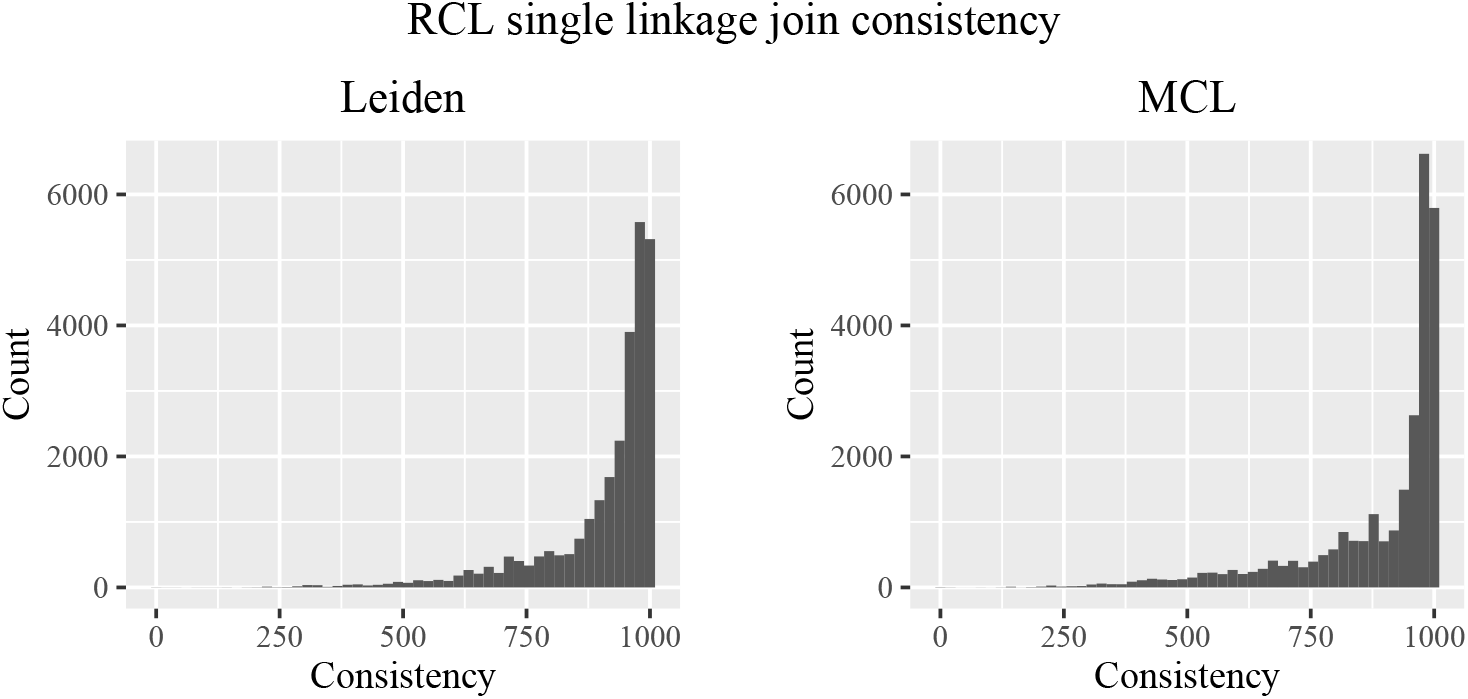
For the two input ensembles Leiden and mcl, the distribution is shown of the consistency value at which nodes are joined in the tree.

The sizes of the subtrees at different consistency levels in the RCL tree are of interest as they reflect the cluster structure at that level. The RCL outcome consists of one or more trees containing a large number of joins, most of which involve a leaf node or a small subtree with few nodes (Figure 9.2). Recall that the smallest size of two subtrees that are merged in a parent node is called the *split* of that parent node (Definition 3.6). For most internal nodes the split is very small. These are complemented by internal nodes that are merges between subtrees of substantial size, representing phase transitions in the RCL tree from one level of granularity or resolution to the next. This is illustrated in Figure 9.3, showing for each internal node *n* both its split and its consistency. The number of nodes with a split at least 100 is 121, with consistency values among these nodes roughly in the range 100 – 750 and a downward consistency value trend for increasing split size.

**Fig. 9.2:**
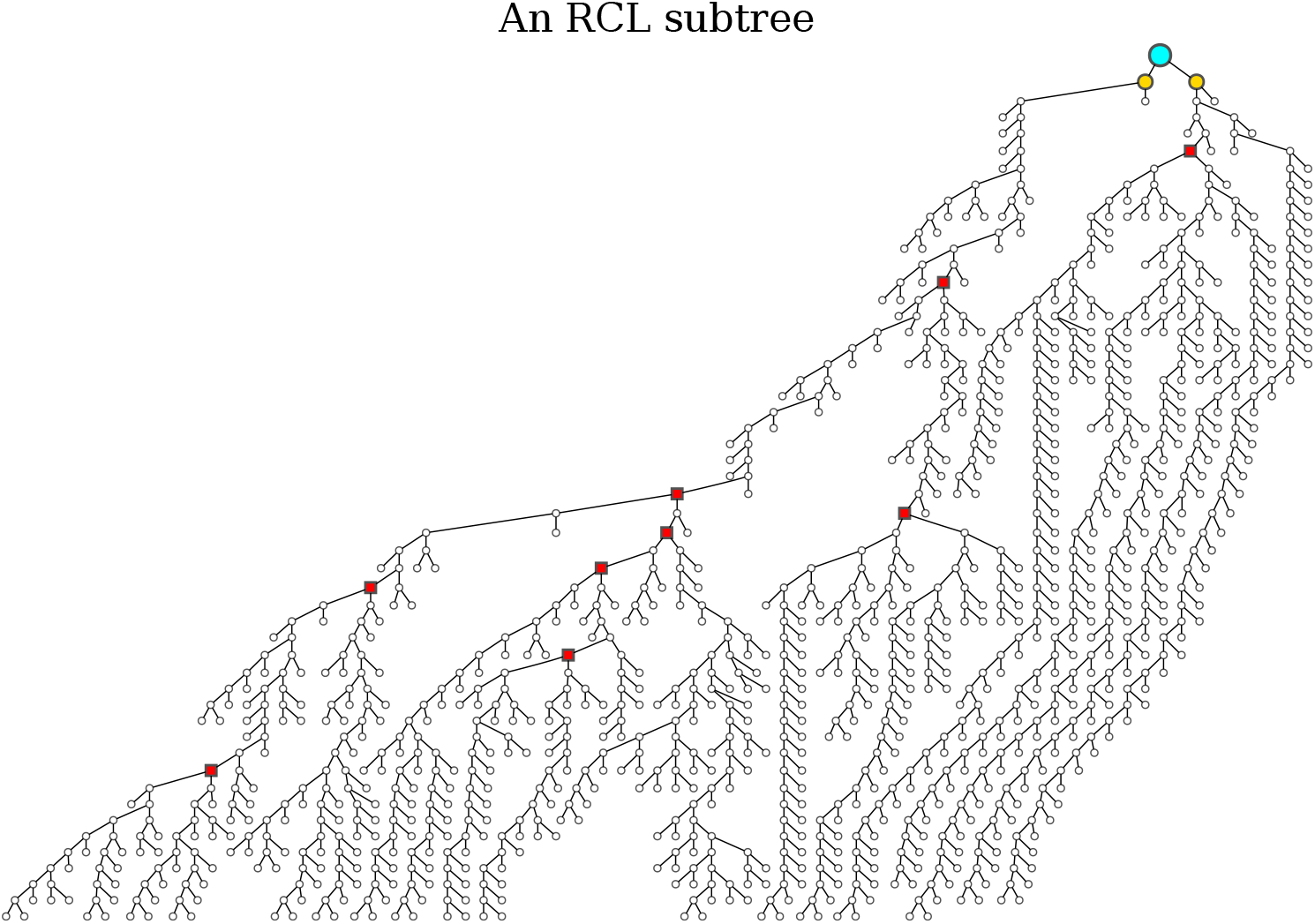
A subtree of the kidney data RCL tree. Branch lengths are uniform and do not represent consistency value differences. The full subtree has 1727 nodes, but it is only shown up to a depth of 52 nodes, on a further reduced set of 1204 nodes. The split of the top node is 329, the size of the smallest of its two subtrees, both marked in gold. Nodes marked in red have a split of at least 50.

**Fig. 9.3:**
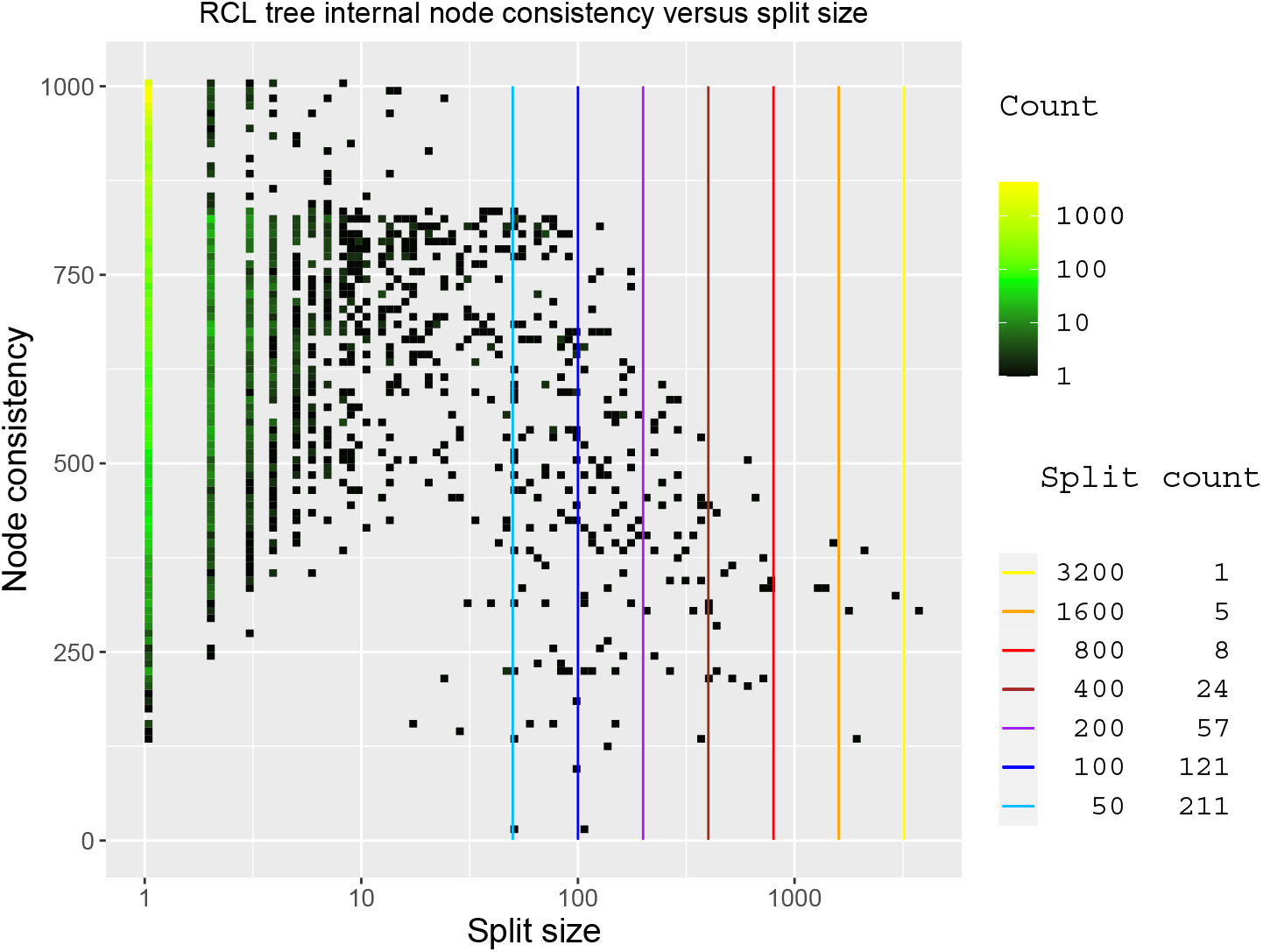
For the RCL tree of the kidney data mcl input ensemble, for each internal node its consistency value versus its split (the smallest of the sizes of the two subtrees below it) is shown with density of counts. The split count table tallies the number of nodes where the smallest of the two sizes is at least *r*, for *r* ∈ {50, 100, 200, 400, 800, 1600, 3200}.

The maximum consistency score 1000 is attained if elements always cluster together and the clusters where this occurs are all mutually consistent. The co-clustering that informs a certain size of clusters is present in input clusterings of similar or coarser granularity. For larger RCL clusters less evidence is available in the input ensemble and the associated consistency level is accordingly lessened, explaining the downward trend observed in Figure 9.3. RCL consistency is affected by two factors, cluster discrepancies and cluster coarseness. I do not attempt to gauge the effect size of these factors here. This is potentially interesting for future analysis, for example to annotate clusters with such information.

The split simplification algorithm (SSA) given in Listing 2 uses some of the nodes with high split size as anchors to derive a multi-resolution view consisting of a small *nested* RCL ensemble of flat clusterings. The goal of the algorithm is to find clusters at different scales, from granular to coarse-grained. This approach differs from the standard approach of cutting the input tree using different similarity level cut-off values. The input ensemble to RCL was chosen such that these differences in scale exist (Figure 8.2), as further illustrated in Figure 9.3.

Below three concepts are singled out to aid the description of SSA.

A node *x* in a binary tree 𝒯 corresponds to a set or cluster, namely all the leaf nodes that are below *x*. A clustering derived from 𝒯 can be represented as a collection of nodes *C* in 𝒯, such that each leaf node *n* encounters exactly one element in *C* in the path from *n* to the root of 𝒯, implying that no node in *C* can be a descendant from another node in *C*. If node *x* is a descendant of node *y*, then the set/cluster associated with *x* is a proper subset of the set/cluster associated with *y*.

Given a resolution size *r*, a node *x* in a binary tree 𝒯 with children *y* and *z* is called *r*-dominant if 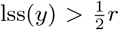 or 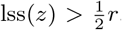, that is, if there is a descendant of *x* that has a split larger than 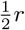.

Given a resolution parameter *r* and a binary tree 𝒯, a flat clustering RES_*r*_ (𝒯) is defined as the smallest set of nodes in 𝒯 such that i) RES_*r*_ (𝒯) is a clustering ii) it contains no *r*-dominant nodes.

Thus, RES_*r*_ (𝒯) is the smallest set of nodes that forms a clustering, such that there is no node contained within it that has a child with a largest sub-split greater than 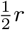. Such a set can be found by descending recursively from the root of the tree *T* (Listing 2). Moreover, if *r < s* then RES_*r*_ (𝒯) is a subclustering of RES_*s*_(𝒯), and RES_*s*_(𝒯) can be used as a starting set of nodes to compute RES_*r*_ (𝒯).

This procedure results in two types of clusters. Consider a node *t* with children *u* and ν. The first type is the node *u* when *t* is *r*-dominant and node *u* was descended towards because 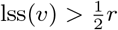, whereas 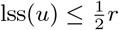. Note that this does not imply any bound for the size of *u*. The second type is the node *t* where 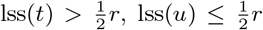, and 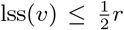, implying that *t* has size greater than *r*, has a split larger than 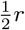, and is itself not *r*-dominant.

A simple algorithm, provided in Listing 2, suffices to compute the clusterings 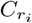 for a range of different *r*_*i*_. A suitable choice for such a range uses a log scale. For single-cell data for example, the analysis in the next section utilised the range of values {100, 200, 400, 800, 1600, 3200}. The smallest value in this scale - call it the resolution limit - is a matter of choice, and will depend on the nature of the data analysed and the granularity of the most fine-grained clusterings among the input ensemble. The question whether the smallest clusters thus detected are, for example, biologically meaningful if the input data is single-cell data, is a matter for the practitioner. After computing a cluster hierarchy with SSA, additional criteria such as marker gene expression can be used to decide which level of granularity and which clusters are of interest.

The benefit of the split simplification algorithm (SSA) is that it finds a nested set of clusters at the different scales provided. If one has reason to believe that rare cell populations may be present, RCL provides an easy way to probe this and will show small clusters embedded hierarchically in larger clusters. In my experience across various datasets SSA results in practice in a well-proportioned hierarchy of clusters exhibiting different scales of grouping, but alternatives to SSA can of course be formulated. Any other method for simplifying the tree will necessarily be consistent. That is, its clusters will either subcluster, supercluster, or be disjoint with the clusters derived by SSA. In light of this I do not view the values in the logarithmic scale used by SSA as RCL parameters, as the RCL tree on which SSA operates is the fundamental result, for which SSA supplies a simplified view. On the other hand tree simplification is necessary to interpret the RCL tree. It is possible that in certain applications a different choice of simplification may yield better results. In that sense SSA is a pragmatic choice where a single best approach is unlikely to exist. In cases where the hierarchical structure within a certain RCL cluster is of very particular interest, it may be worthwhile to examine more closely the branching behaviour of the RCL tree below the tree node corresponding to this cluster.

## 10. RCL data representation and outputs

The RCL reference implementation uses the split simplification algorithm (SSA) and thus generates a small set of flat clusterings from the tree that results from applying single linkage clustering to the RCL matrix.^7^ As the split simplification algorithm descends the tree as the resolution size is decreased, the corresponding flat clusterings generally will have a proportion of clusters that are singletons or of small size.

The reference implementation uses a simple notion, namely that any cluster smaller than the resolution limit (i.e. the smallest resolution size provided) is referred to as a *residual cluster*, and an element within such a cluster is called a *residual element*.

It is up to the user of the software to decide the smallest size of cluster that is possibly of interest, and such a decision may be informed by the nature of the experimental data. It bears emphasizing that a node may be part of a proper (non-residual) cluster *C* at one resolution size, but becomes part of a residual cluster as *C* splits into smaller clusters at a lower resolution size (see the example below). For large resolution sizes there will be very few residual elements; as the resolution size is decreased and smaller clusters are retrieved, the tree descent mechanism will increase the number of clusters and the number of residual clusters and elements will increase accordingly.

### Listing 2

RCL tree simplication algorithm.

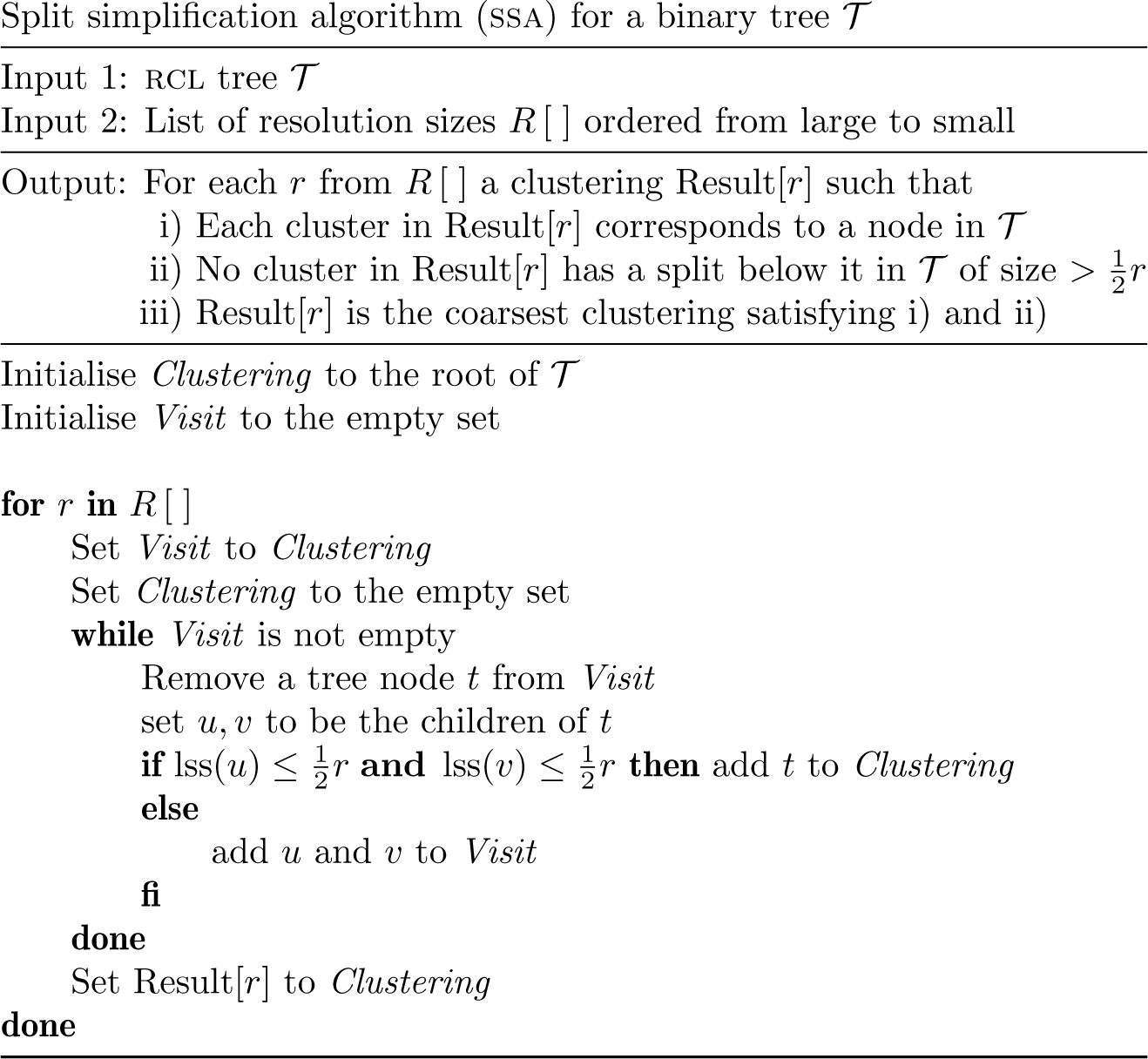

For the flat clusterings derived by SSA the notion of residual element has no further consequence. The reference RCL implementation (Section 13) additionally provides a tabular representation where all the flat clusterings are arranged into a hierarchy. In order to make this hierarchy manageable and accessible residual elements are grouped together. This is illustrated below with an excerpt from such an output.

| level | type | joinval | N1 | N2 | nesting | elements |
| --- | --- | --- | --- | --- | --- | --- |
| 1 | residual | 0 | 27203 | 2134 | A | 1 5 6 7 8 9 11 13 15 16 17 18 ... |
| 2 | residual | 131 | 6507 | 835 | B_A | 20357 20361 20362 20363 ... |
| 3 | residual | 332 | 3363 | 3 | B_B_A | 22589 23360 26840 |
| 4 | residual | 335 | 1436 | 398 | B_B_B_A | 20579 20700 20767 20768 ... |
| 4 | cls | 535 | 1436 | 1038 | B_B_B_B | 20371 20377 20380 20441 ... |
| 4 | residual | 366 | 1221 | 20 | B_B_C_A | 20617 22318 22377 22601 ... |
| 4 | cls | 407 | 1221 | 695 | B_B_C_B | 20704 20760 20863 20873 ... |
| 4 | cls | 542 | 1221 | 506 | B_B_C_C | 20479 20847 20972 21446 ... |
| 4 | residual | 343 | 703 | 46 | B_B_D_A | 20762 20835 21239 21510 ... |
| 4 | cls | 584 | 703 | 657 | B_B_D_B | 20463 20771 20791 20912 ... |
| 3 | residual | 152 | 1886 | 1093 | B_C_A | 123 9675 11529 15097 17428 ... |
| 3 | cls | 451 | 1886 | 793 | B_C_B | 20369 20477 20490 20498 ... |
| 2 | cls | 294 | 6507 | 423 | B_D | 20391 20404 20470 20478 ... |
| 2 | residual | 382 | 5350 | 18 | C_A | 10149 10642 11076 11432 ... |
| 3 | residual | 399 | 3205 | 10 | C_B_A | 1951 10094 11554 12511 ... |
| 4 | residual | 407 | 1697 | 1039 | C_B_B_A | 348 470 1520 1924 2457 ... |
| 4 | cls | 605 | 1697 | 658 | C_B_B_B | 9638 9641 9643 9646 9653 ... |
| 4 | residual | 456 | 1498 | 8 | C_B_C_A | 250 2390 9987 10159 11534 ... |
| 4 | cls | 513 | 1498 | 807 | C_B_C_B | 173 407 417 475 2271 2324 ... |
| 4 | cls | 480 | 1498 | 683 | C_B_C_C | 2228 9575 9595 9599 9606 ... |
| ... |  |  |  |  |  |  |
| .. |  |  |  |  |  |  |
| . |  |  |  |  |  |  |

The table is ordered such that cluster structure at all levels is represented by contiguous rows in the table, as indicated by the `nesting` column. Using this ordering in a cluster/gene heatmap thus realises the RCL hierarchy in the heatmap (Figures 11.1-11.2) without further need for clustering and is made explicit by additionally supplying a dendrogram.

**Fig. 11.1:**
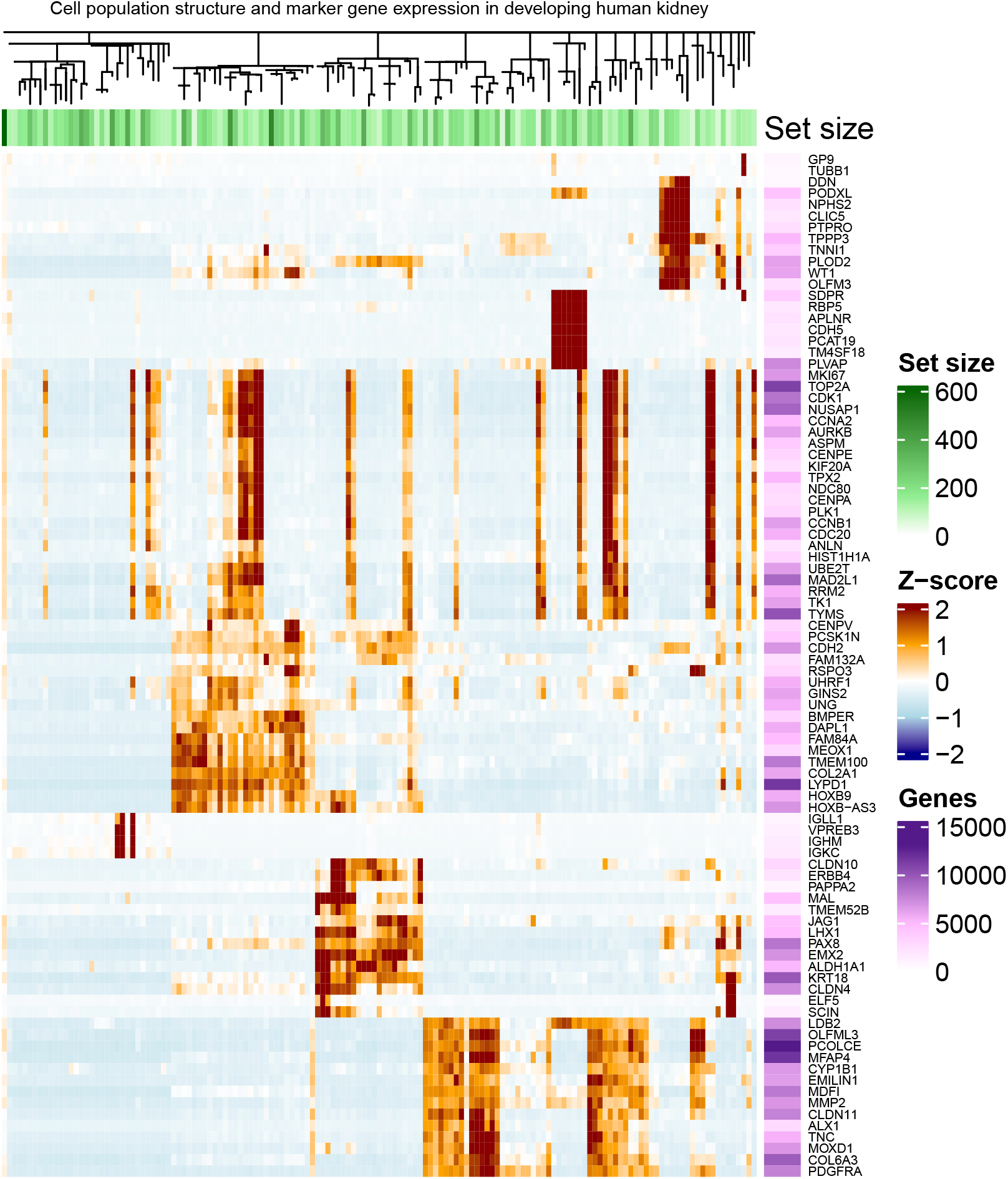
ComplexHeatmap [37] of marker genes for RCL clustering of cell transcriptomes from 27203 developing kidney cells. RCL clustering was performed as described in Section 8. Heatmap columns correspond to the rows in the tabular representation of the most fine-grained clusters (Section 10). Coarser levels are visible in the column clustering and the associated dendrogram. Clusters of residual nodes (Section 10) can be recognised as horizontal bars without a vertical descent. (Heatmap continued on the following page).

**Fig. 11.2:**
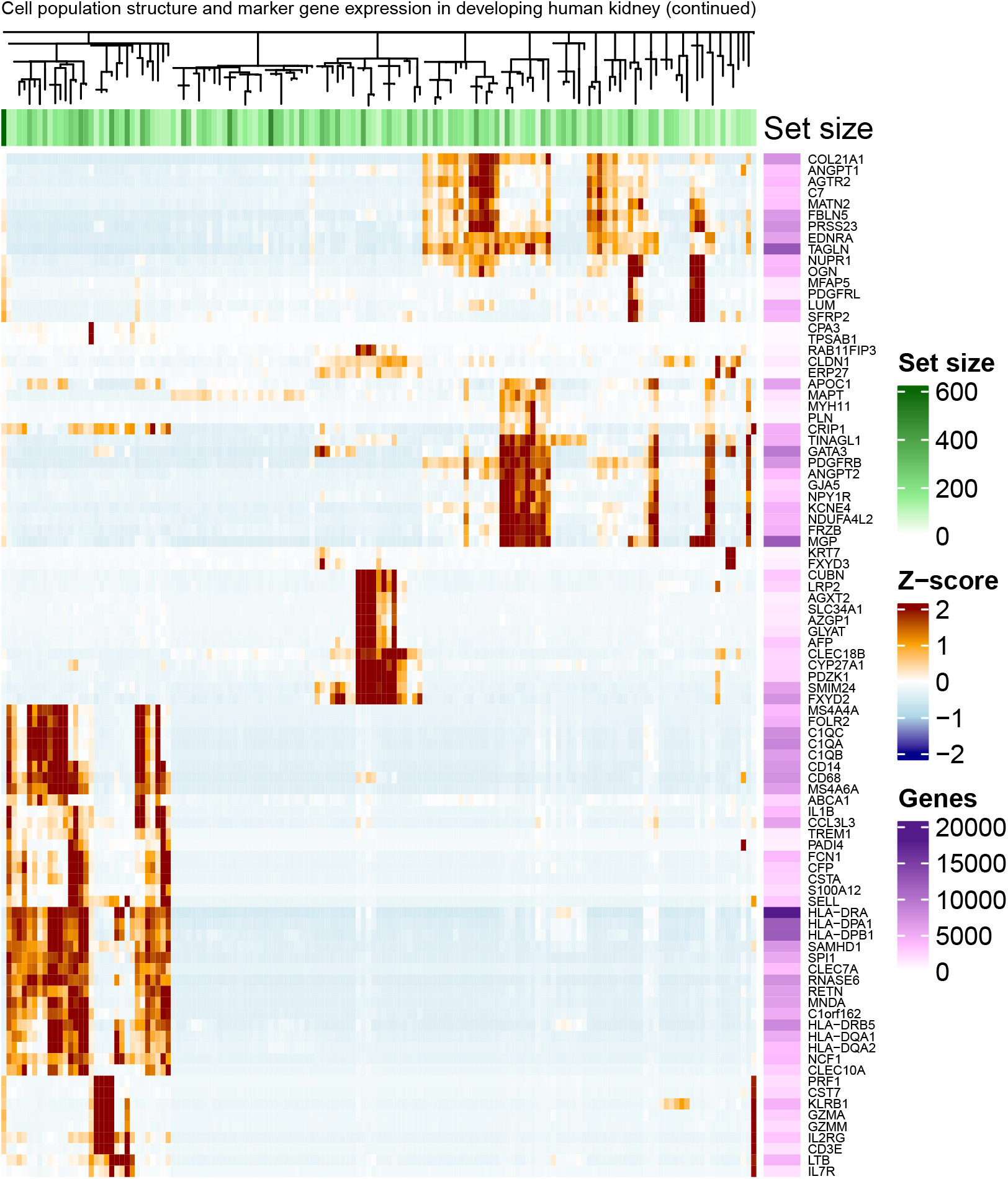
Continuation of the heatmap of marker genes for RCL clusters across developing kidney cell transcriptomes in Figure 11.1.

In this example RCL was run on the developmental kidney data with resolution values 400, 800, 1600, 3200, thus the resolution limit was considered to be 400. For the purpose of this example it was set to a relatively high value; a recommended set of standard values is 100, 200, 400, 800, 1600, 3200. Each data row represents a cluster at the most fine-grained level from the RCL flat clusterings. The `N2` column indicates the size of the cluster, the N1 column indicates the size of the parent cluster; the nesting column encodes the `nesting` structure of the clusters. The `elements` column (here generally shown truncated) contains all identifiers for the elements of a cluster. To obtain a coarser cluster one combines these elements, so that, for example, to obtain the elements for the cluster `B_B` one takes all rows where the `nesting` identifier has `B_B` as a prefix.^8^ The reference implementation provides software for such manipulations, as well as for translating element identifiers to meaningful labels. In the example there are eight rows combining to a total of 3363 elements for the `B_B` cluster.

All nesting identifiers that have suffix `A` represent a set of residual clusters *joined together*. Thus, the first set labelled `A` comprises 2134 elements that reside in clusters of size smaller than 400 even at the coarsest resolution limit of 3200. This number is unrealistically large in this example due to the high resolution limit. If the resolution limit is set to 100 as recommended, the number of residual elements for the *A* set is reduced to 611. The parent clustering of *A* is the root clustering of all 27203 elements (cells) in the data set. This top-level residual cluster *A* represents volatile nodes that do not participate in a cluster of sufficient interest/size at any level of consistency, having co-clustered ambiguously across the input ensemble.

The *B* cluster has 6507 elements and is represented by several immediate subclusters. The first of these is the residual set `B_A`, consisting of 835 elements. These elements are first-class citizens of *B*, but will generally have joined late and are not part of a sufficiently large cluster at any lower resolution level. The elements in `B_A` inherently belong to *B* but cannot be assumed to have a distinct separate identity as they collate separate small subtrees in the RCL tree below the node that represents *B*. The 3363 elements in `B_B` on the other hand do correspond to a node and subtree in the RCL tree and thus group together naturally. Their grouping may be (biologically) meaningful in a narrower sense beyond their membership of B. By extension these considerations apply to all clusters at all levels.

For single-cell data the reference implementation provides software to visualise the tabular representation described here in a heatmap of marker gene expression across RCL clusters, an example of which is given next.

## 11. RCL multi-resolution clustering of developing kidney cells

Although they remain the default way to represent and interrogate single-cell transcriptomic data, non-linear dimension reduction techniques such as umap [34] or t-SNE [35] represent a limited [36] snapshot of single-cell transcriptomes. Heatmaps offer an alternative visualization approach, not limited by the issues of over-plotting that plague scatterplots [37]. However, for heatmaps to provide a useful alternative for exploration of scRNA data, cells must be clustered and ordered in a biologically meaningful way. mcl or Leiden clusterings organized into a tree structure by RCL can potentially solve this problem.

To test this I applied RCL to single-cell transcriptomes from the developing human kidney [13], using the RCL results for the mcl input ensemble described in Section 8. Resolution sizes were set to 100, 200, 400, 800, 1600 and 3200. RCL clusters were collated in a simplified hierarchy as described in the previous section, so that for any given cluster all its immediate subclusters below the smallest resolution size 100 were gathered in a single residual subcluster. The clusters were ordered such that the coarser clusterings at all higher resolution levels were preserved, as illustrated in the example output in the previous section. Marker gene analysis was applied for each cluster at the most fine-grained level separately, using the `quickMarkers` function from the SoupX software [38]. The top two marker genes for each cluster were selected provided that their tf-idf value was at least one, resulting in a combined set of 181 marker genes. These genes have widely varying roles in cell function due to the large range of fine-grained clusters from which they are derived.

To interrogate cluster identity and cell types a heatmap for these clusters and marker genes was created using the ComplexHeatmap software [37]. Columns and rows in the heatmap represent clusters and marker genes, respectively. To evaluate the biological meaning of the resulting clustering and hierarchy, I plotted the (row normalized) average expression in each cluster for each marker gene (Figures 11.1 and 11.2). The heatmap column ordering is fully pre-defined by the RCL ordering of clusters as described in the previous section. Additionally the RCL software encodes the multi-resolution hierarchy in Newick format, using the consistency levels at which clusters form to define branch lengths. This format is accepted by ComplexHeatmap and rendered as a dendrogram showing the RCL hierarchy. The branch length of a residual cluster relative to its parent cluster is set to zero, so that residual clusters (the leftmost cluster in any subtree) have no vertical descent. Residual clusters of size below 50 were excluded. The heatmap row ordering is defined by ComplexHeatmap itself based on clustering of normalised marker gene expression.

The leftmost column in the heatmap represents the top-level residual cluster with 611 elements, that gathers clusters of size below the resolution limit 100 as described in the previous section. These clusters are associated with very low consistency values and thus with nodes that are volatile and co-cluster poorly. A residual cluster is formed, at any level, as a gathering of small subclusters of a parent cluster. Hence gene expression in the residual cluster may (only) be assumed to be within the representative range for this parent cluster. For the top-level residual cluster the parent cluster is the entire dataset, and thus it can be expected to exhibit expression across a wide range of marker genes. This is indeed the case, as can be observed by scanning the leftmost column.

The top split in the hierarchy (Figures 11.1 and 11.2) clearly separates leukocytes (left-most clade) from all other cells. Other major splits clearly represented endothelial cells (marked by CDH5 and PCAT19 expression), intermediate/proximal tubular cells (PAX8, CUBN expression), mesenchymal/interstitial cells (PDGFRA, MMP2), cap mesenchyme cells (MEOX1, TMEM100), and podocytes (PODXL, PTPRO). Furthermore, within each clade smaller sub-populations were found representing cell-type-specific cycling cells (expression of MKI67, TOP2A). This segregation of cycling cells across clades is a distinct advantage over umap/t-SNE, where cycling cells often group together regardless of cell type.

To further investigate if the fine splits on the tree were biologically meaningful, I identified clusters with marker genes not shared by any other cluster, considering the top five marker genes for each cluster (Figures 11.1 and 11.2 only show the top two). This revealed rare, but biologically indisputable populations of cells including Mast cells (TPSAB1, TPSB2 expression, 203 cells), plasma cells (IGKC, JCHAIN, 208 cells), and a specific sub-population of interstitial cells (NTRK3, MYH11, 167 cells). Expression of four interstitial marker genes is shown in Figure 11.3 in a umap plot of all 27203 developing kidney cells. This expression overlaps to a large extent with the sub-population detected by RCL, shown in Figure 11.4 embedded in larger scale cluster structure as found by RCL. This is local to a region of the umap plot that was found to be challenging to annotate [13]; the interstitial cells are embedded in a larger cluster of mesenchymal cells, that likely give rise to a range of extraglomerular cell types but lack well established marker genes that can be used to definitively annotate them. Taken together, these findings show that the tree structure and clustering produced by RCL is biologically meaningful and can reveal rare, biologically meaningful, sub-populations of cells that are easily missed with current approaches.

**Fig. 11.3:**
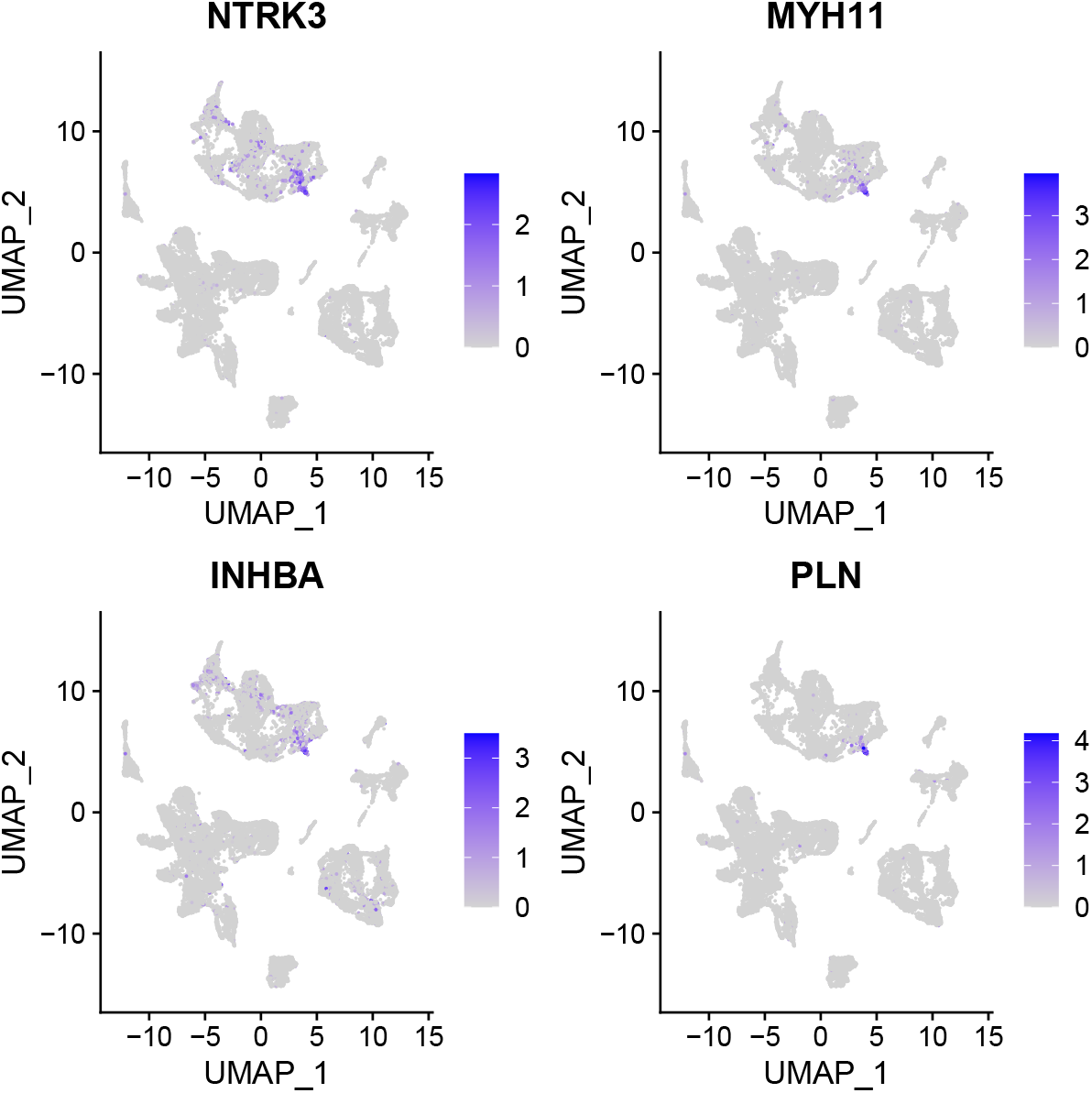
Expression of interstitial cell marker genes, largely localised to the small cluster of RCL cells shown in Figure 11.4.

**Fig. 11.4:**
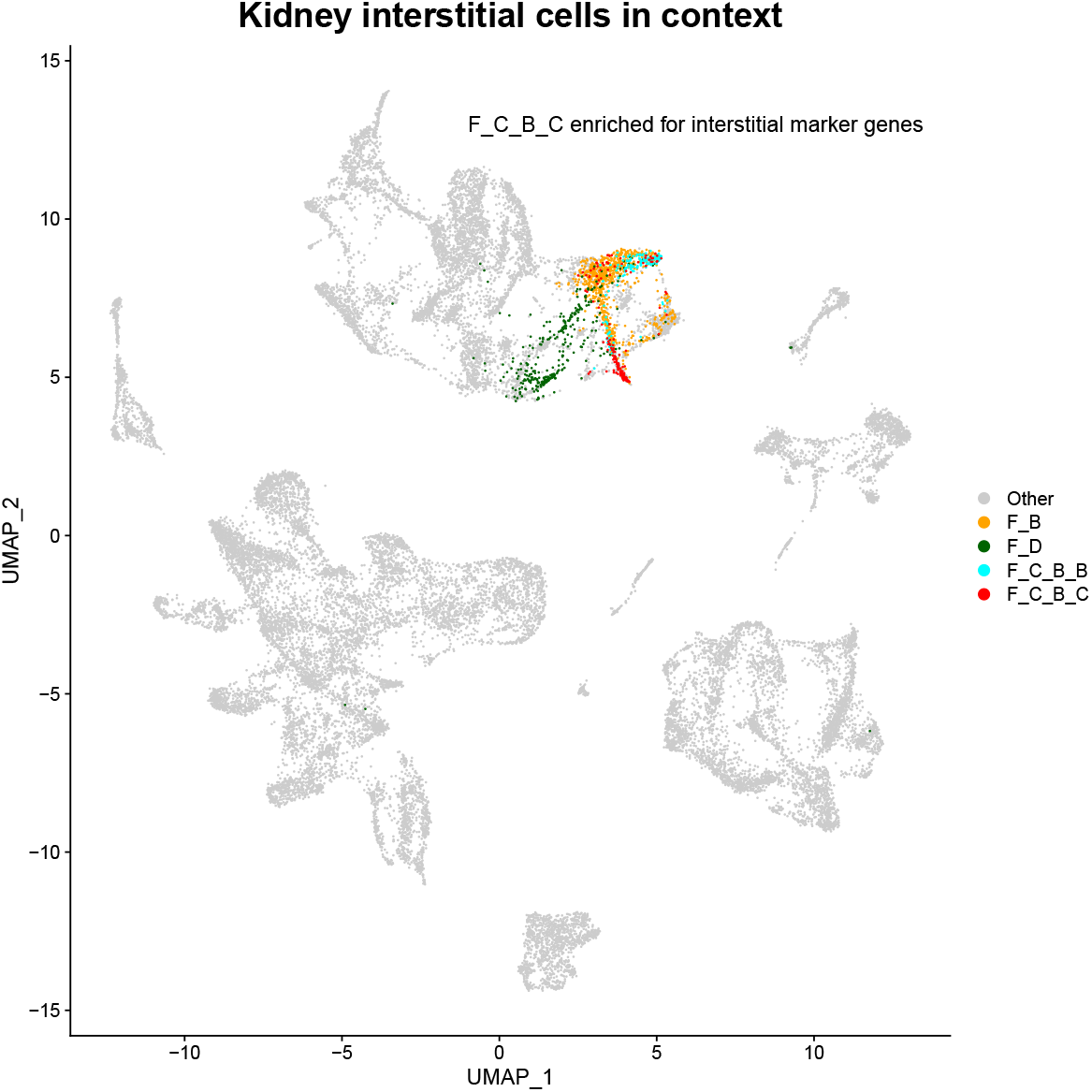
The RCL cluster capturing interstitial cells (`F_C_B_C`). It has 167 cells, enveloped by successively larger clusters of 356 cells (`F_C_B`) and 1727 cells (`F`).

## 12. Complexity, speed and implementation

Computation of the RCL matrix is of order 𝒪(*m*(*p*^2^ + log(*m*))) where *m* is the number of edges in the network and *p* is the number of clusterings. Computation of the RCL network dominates the running time of the implementation described here and is of order 𝒪(*mp*^2^). Computation of the single linkage hierarchy is of order 𝒪(*m* log(*m*)). Although potentially more computationally demanding due to the logarithmic factor, for the data tested (with *d* = 27k elements, 2.3M edges and *p* = 24 clusterings) this step was much faster. The factor *m* log(*m*) is due to sorting of edges based on weight. In the setting of networks, where all pairwise similarities of interest have already been computed, sorting and processing all edges is simple and effective. Sorting algorithms are highly optimised with low overhead, that is, constant factors in the computational cost are small. After sorting, the edges are processed to construct the single linkage hierarchy, using linked lists to efficiently update the correspondence from leaf nodes to internal root nodes.

For computation of the RCL network the factor *p*^2^ is clear-cut and derives from pairwise comparisons for all partition combinations. As before the number of nodes or data elements is denoted by *d*. For each comparison of partitions we compute the contingency matrix *T* with *T*_*ij*_ = | gcs(𝒫_*i*_, 𝒫_*j*_)|. Partitions can be stored as a sparse matrix with *d* elements; the contingency matrix is obtained by multiplying two such matrices, with a cost in 𝒪 (*d*). This cost will be dominated by the next step however. For each set in gcs(𝒫_*i*_, 𝒫_*j*_) (an intersection of two clusters in 𝒫_*i*_ and 𝒫_*j*_ respectively) we visit its elements and update a subset of its neighbours in the RCL network. Since each node is part of just one intersection, this means we update each edge at most once, leading to an overall complexity of 𝒯 (*mp*^2^).

On a dataset with *N* = 27k elements and *p* = 24 clusterings the method takes less than a minute. An RCL implementation is available as part of the mcl sparse matrix and clustering software, as of version 22-282 (https://github.com/micans/mcl).

## 13. Discussion

I have introduced a novel consensus clustering method; Restricted Contingency Linkage or RCL is a fast and parameter-free method to integrate and reconcile a set of flat clusterings, subsuming any method that produces such clusterings. For widely used methods such as Leiden and mcl this opens up the possibility to utilise cluster structure at different levels of granularity. This obviates the search for a best resolution parameter or inflation parameter, where any specific such value both is a compromise across the entire dataset and fixes the scale of the view. As such RCL offers an alternative to hierarchical clustering methods that are well-established in the literature, but are rarely used in practice for the purpose of identifying and relating cluster structure at different scales. The fundamental advance of RCL is the matrix object that encodes the co-clustering relationships in the input clusterings. This RCL matrix is a considerable improvement on a similar object, the association matrix, that is part of the prevailing paradigm in consensus clustering. I have demonstrated how the entries in this matrix are richly differentiated (Sections 4-8), encoding the strength and consistency of co-clustering at different scales. Single linkage clustering of the matrix accurately reflects these different scales whilst equally accounting for the discrepancies amongst the input clusterings.

A fast and efficient command line implementation of RCL is available as part of the mcl software (https://github.com/micans/mcl/RCL). All the results and visualisations presented here for the developmental kidney data can be created using simple short recipes, available as part of the documentation. In the case of single-cell data the starting point for these recipes is a set of three data files, the raw expression data in matrix market format, the gene names, and the barcode/cell names. The tabular output format described in Section 10, by default produced by RCL, is suggested as the easiest data format to load results.

Current practices in application areas such as single-cell and protein sequence analysis are geared towards flat clusterings. Hierarchical data brings a larger burden for data administration and visualisation, even more so as data sets grow in size and complexity. These matters of presentation are outside the realm of the core RCL method for deriving a multi-resolution data description. Presentation and navigation of large-scale data is likely to remain an area of research and development for the foreseeable future. Hierarchy-informed variants of umap exist, such as humap [39]. I propose that it is beneficial to separate data classification from presentation. Hence it will be desirable that such methods can be adapted to allow an externally-computed hierarchy as input. A second issue is that visualisation techniques that create a low-dimensional spatial embedding (such as umap and t-SNE) will struggle increasingly as data sizes increase. The more abstract representation of the heatmap for marker gene expression across a hierarchical clustering (Section 12) is a useful and scalable complementary alternative. It can serve as a comprehensive first point-of-call static chart of the data, mapping its salient features from detailed to abstract and indicating populations of interest at different scales, from small and specific cell types to large unifying clades.

It is a priori not known which clusters generated by RCL are biologically meaningful. The clusters at the most granular levels may be overclustered, that is, represent a grouping in the data that has arisen by chance or as a sampling artefact. Nevertheless, a small cluster may be meaningful and represent for example a rare cell type or cell state population. Probing such meaning and identifying interesting clusters requires additional analyses, for example by applying tests for marker genes, as in Section 8. The RCL software enables such marker gene analysis in a single command. Additionally the resolution list provided to RCL can be adapted to change the level of detail observed. This is a separate step in an RCL analysis that is fast and easy to re-run. In single-cell data analysis other approaches exist that incorporate significance testing of gene expression across clusters [40, 41]. By taking into account variation of expression at the level of individual genes they allow more direct evaluation and control, thus aiming to avoid overclustering. These methods have stronger modelling assumptions and depend on a more complex inference machinery. The choice between such a method and a purely abstract clustering approach like RCL will depend on application area, as well as ease, speed and availability of software.

An intrinsic part of RCL is the notion that some data elements are less equal than other data elements, in that they cluster inconclusively and switch allegiance between different sets of nodes in different clusterings, corresponding to discrepancies between those clusterings. In the case of clusterings of varying granularity/scale, this points to a data landscape where the ridges that separate clusters differ depending on the vantage point. An area of future study is to investigate these dynamic ridges by considering the consistency measure and how it is affected by clustering discrepancies and variation in coarseness respectively. One aim is to achieve a more controlled and principled way of deriving for example shared nearest neighbour networks from high-dimensional data.

Although RCL is parameter-free, the choice of input ensemble (Section 8) still merits attention. A question not addressed here is whether it is beneficial to have larger input ensembles with more frequently sampled resolution or inflation parameters. A possible addition to RCL may then be to allow reduction of the number of pairwise comparisons by diagonal banding, that is, rather than performing all pairwise comparisons on the input ensemble, to just compare clusterings that are somewhat near in terms of cluster granularity, for example by setting a bound on the discrepancies between clusterings (Section 3) beyond which they are not compared.

Cluster analysis is a foundational pillar of data exploration. RCL adds exploration along the dimension of scale, with the measure of consistency quantifying the cohesiveness of the groupings. Visualisation of clustering outcomes at different scales of resolution, adorned with consistency, will allow analysts to engage with their data in a more informed and intuitive way.

## 14. Acknowledgements

Matthew Young engaged in many helpful discussions, suggested the use of (quick)marker genes and (complex)heatmaps to summarise and visualise single-cell data analyses, provided the kidney data set and analysed the application of RCL to this data. Vladimir Kiselev encouraged me for a long time to test mcl with single-cell data - it finally happened. Simon Murray ran Seurat analyses and clusterings, and provided data sets for testing. Alexander Predeus suggested data sets and possible uses for RCL. This research is the current arrowhead of a long arc of clustering in bioinformatics in which Anton Enright has been hugely influential. Izaak van Dongen, Lena van Dongen and Leopold Parts suggested many improvements to the manuscript. I am very grateful for all their help and encouragement.

## Footnotes

1 The deficit is exactly equal to the number of components at the coarsest level of the co-clustering relationship.

2 Axiomatic analyses exist as well, wherein *a consensus classification is required to satisfy a set of axioms*, and researchers investigate *the existence and uniqueness of solutions* [16].

3 Discrepancy is closely related to the split-join distance between partitions introduced in [19]. This split-join distance, denoted sjd, is a metric distance on the space of partitions of 1, …, *d* and is defined as 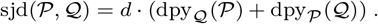 The split-join distance thus counts the number of elements that need to be moved via splits and joins to transform 𝒫 into 𝒬 or vice versa.

4 Colours were reused between the different partitions; this is immaterial for the purpose of clustering, that is, there is no intrinsic meaning associated with clusters in different 𝒫_*i*_ having the same colour.

5 This is implicitly arranged by the set-finding function **S** taking the value of the empty set if *x* and *y* do not co-cluster, and scy being zero if any of its arguments is the empty set.

6 That is, counter-examples may exist but are likely to be tortuous and artificial, as it would require two nodes not connected in *G* to co-cluster directly before being joined transitively by co-clustering of the nodes on a path between them.

7 These are produced in different formats, including the common two-column format used by Seurat that has a *label* and its *cluster ID* in each line.

8 Equivalently one can take the first such cluster `B_B_A` and all subsequent rows for which the `level` value is larger than that of `B_B_A`.

